# Screening the human miRNA interactome reveals coordinated upregulation in melanoma, adding bidirectional regulation to miRNA networks

**DOI:** 10.1101/2024.05.10.593635

**Authors:** Faezeh Jame-Chenarboo, Joseph N. Reyes, Nicholas M. Twells, Hoi-Hei Ng, Dawn Macdonald, Eva Hernando, Lara K. Mahal

## Abstract

Cellular protein expression is coordinated post-transcriptionally by an intricate regulatory network. The current presumption is that miRNA work by repression of functionally related targets within a system. In recent work, upregulation of protein expression via direct interactions of mRNA with miRNA has been found in dividing cells, providing an additional mechanism of regulation. Herein, we demonstrate coordinated upregulation of functionally-coupled proteins by miRNA. We focused on CD98hc, the heavy chain of the amino acid transporter LAT1, and α-2,3-sialyltransferases ST3GAL1 and ST3GAL2, which are critical for CD98hc stability in melanoma. Profiling miRNA regulation using our high-throughput miRFluR assay, we identified miRNA that upregulated expression of both CD98hc and either ST3GAL1 or ST3GAL2. These co-upregulating miRNAs were enriched in melanoma datasets associated with transformation and progression. Our findings add co-upregulation by miRNA into miRNA regulatory networks and adds a new bidirectional twist to the impact miRNA have on protein regulation and glycosylation.

## Introduction

Cells are defined by protein expression patterns that require precise regulation^1,2^. microRNA (miRNA, miR), small non-coding RNA, provide such precision^3,4^. The canonical view of miRNA action is that miRNA dampen protein expression through binding the 3’untranslated region (3’UTR) of a transcript (down-miR)^3^. Recent work from our laboratory has challenged this view, demonstrating that miRNA can also increase protein expression through direct interactions with the 3’UTR within actively dividing cells (up-miR)^5,6^. Our work was in concordance with previous reports of enhanced translation via miRNA:mRNA interactions in quiescent cells^7,8^. Argonaute 2 (AGO2), a fundamental part of the miRNA machinery, is required for upregulation by miRNA. In contrast, TNRC6A (GW182), which is a crucial component of the complexes required for downregulation, was not required for upregulation – providing strong evidence that up-miRs do not work by a “relief of repression” mechanism^6,9,10^. Given the important role miRNA have in regulating levels of proteins that coordinate specific biological functions, we would expect that upregulatory interactions would act cooperatively to tune protein expression in these regulatory networks. Herein, we show incorporation of upregulation into such networks, providing evidence that miRNA co-upregulate related targets and positioning them as bidirectional tuners of expression.

In a recent study, we identified a functional relationship between the CD98 heavy chain (CD98hc) and the *α*-2,3-sialyltransferases ST3GAL1 and ST3GAL2 in melanoma^11^ (Fig. 1a). CD98hc, encoded by *slc3a2*, is a type II transmembrane glycoprotein ubiquitously expressed at high levels on many cell types. It is a part of the neutral amino acid transport complex LAT-1^12,13^. CD98hc has roles in both physiological and pathological contexts, including in B- and T-cell expansion, integrin signaling and cell proliferation^12,14-16^. It is a target of anti-cancer antibodies in current clinical trials due to its importance in the biology of various solid tumors (e.g., bladder^17^, breast^18^, lung^19^, melanoma^11,20^) and hematological cancers^15,21^. In melanoma, *α*-2,3-sialylation of CD98hc by the enzymes ST3GAL1 and ST3GAL2 was found to be important in the stability of CD98hc, protecting it from proteasomal degradation^11^.

**Fig. 1.**
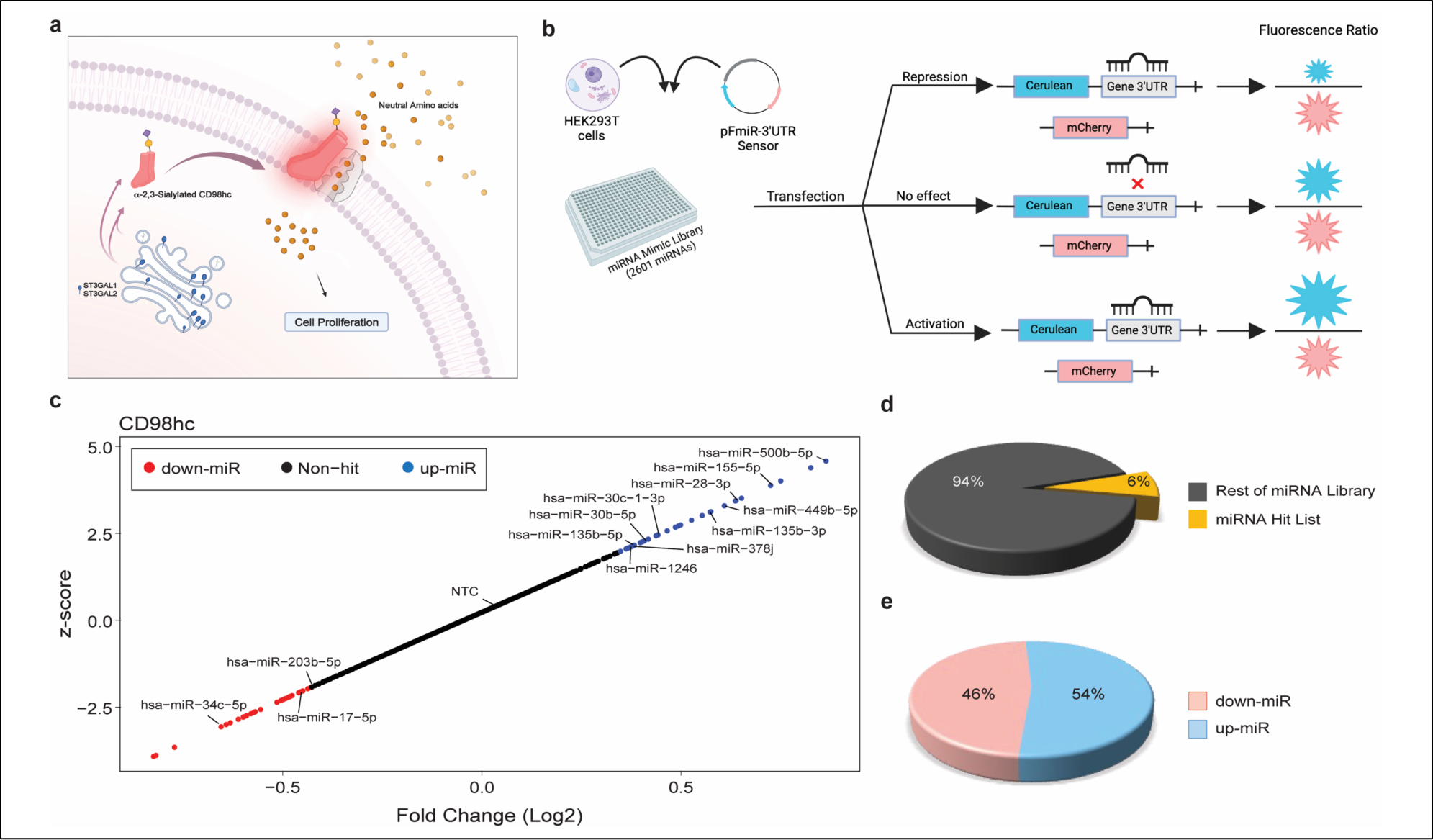
Comprehensive map of CD98hc regulation by miRNA. **a.** α-2,3-Sialylation of CD98hc by ST3GAL1 and ST3GAL2 stabilizes this essential protein in melanoma^11^. **b.** miRFluR assay: 3’UTR of gene of interest (CD98hc) is cloned downstream of Cerulean in pFmiR construct (pFmiR-CD98hc). Plasmid is then screened with human miRNAome library (Dharmacon, miRbase v. 21) in HEK293T cells. Ratio of Cerulean: mCherry signal normalized by median ratio indicates miRNA impact. **c.** Scatterplot of miRFluR data for 3’UTR of CD98hc. miRNA in the 95% confidence interval by z-score are indicated (down-miRs: red, up-miRs: blue) and validated miRNAs are shown. **d.** Pie chart showing % miRNA hits compared to total library post-QC. **e.** Pie chart indicates % down-miR (48%) vs. up-miR (52%) in CD98hc hit list.

ST3GAL1 and ST3GAL2 transfer sialic acid onto the 3-position of terminal galactose predominantly on *O*-glycans and glycolipids^22,23^. ST3GAL1, which can biosynthesize the tumor antigen sialyl T (Neu5Acα2-3Galβ1-3GalNAc)^24^ plays functional roles in the biology of several cancers including breast cancer^25^, ovarian cancer^26^, and melanoma^11,27^. It is also an important regulator of cell migration^23^ and recently was found to mediate sequestration of CAR-T cells from circulation, inhibiting the cell’s ability to home to cancer tissues^28^. In contrast to ST3GAL1, less is known of the function of ST3GAL2. It is associated with transformation in colorectal cancer^29^ and is highly expressed in pancreatic cancer compared to normal tissue^30^. We have recently shown that it, along with ST3GAL1 and CD98hc, is an essential gene in melanoma^11^.

We posited that, given the functional relationship between CD98hc and both ST3GAL1 and ST3GAL2, that these genes might be co-regulated by miRNA. We screened the human miRNAome for regulatory relationships for all three genes using our high-throughput miRFluR assay. In previous work, we have used this assay to uncover miRNA regulation of the glycosylation enzymes B3GLCT^5^, ST6GAL1 and ST6GAL2^6^, leading to the discovery that miRNA can upregulate protein expression through direct interactions with the 3’UTR of mRNA (upregulatory miRNA or up-miR). Analysis of CD98hc, ST3GAL1 and ST3GAL2 herein provided a regulatory map of these genes, identifying 10 co-up-miRs: 6 between CD98hc and ST3GAL1 and 4 between CD98hc and ST3GAL2. The majority of these co-upmiRs were associated with either melanoma metastasis or progression^31,32^, positioning upregulation as a central part of miRNA regulatory network.

## Results

### Mapping miRNA regulation of CD98hc identifies up- and downregulatory interactions

To map the miRNA regulatory landscape of CD98hc, we utilized our miRFluR assay^5^ (Fig. 1b). Our assay uses a genetically-encoded fluorescent sensor (pFmiR) that contains the GFP variant Cerulean regulated by the 3’UTR of the gene transcript of interest with mCherry as an internal control^5^. Downregulatory miRNA (down-miRs) result in loss of Cerulean (blue), while upregulatory miRNA (up-miRs) cause enhanced protein expression^6^. In our previous work, we analyzed the miRNA regulation of glycosylation enzymes, which are moderate to low abundance proteins^5,6^. In contrast, CD98hc is highly expressed at both the protein and transcript levels in many cancer (e.g. melanoma^20^) and non-cancer (e.g. immune) cells^15,21^. To assess regulation, we cloned the most prevalent 3’UTR for CD98hc into our sensor to create pFmiR-CD98hc (Supplementary Fig. 1). While the 3’UTRs of other genes studied in our system were > 1 kilobase in length; the 3’UTR of CD98hc is only 190 nucleotides (nt). We co-transfected our pFmiR-CD98hc sensor with a miRNA mimic library (Dharmacon, v. 21, 2601 miRs, 50 nM, arrayed in 384-well plates) into HEK-293T cells. All miRNA were represented in triplicate and fluorescent signals were analyzed 48 hours post-transfection. Upon data analysis, we found that the “non-targeting” controls (NTCs) provided with the library were significantly shifted from the median signals within the plates, arguing that they impacted sensor expression. These “controls” are miRNA from other organisms and may have binding sites within human 3’UTRs. Therefore, we median normalized our data (Fig. 1c). After quality control (QC), we log_2_ transformed the data (954 miRs) and then applied a z-score threshold at the 95% confidence interval to identify hits. Our high-throughput analysis found approximately even numbers of downregulatory (down-miRs, n=30), and upregulatory (up-miRs, n=35) miRNA, representing 6% of miRNA interactions passing QC (Fig. 1c-e, Supplementary Fig. 2, Dataset 1).

### CD98hc expression is both up- and downregulated by miRNA in cancer cells

CD98hc is an emerging cancer target, with a therapeutic antibody in Phase I clinical trial^16,18,33,34^. In solid tumors, expression of CD98hc is highly correlated with progression and metastasis^16^. For initial validation, we chose a subset of miRNAs previously shown to regulate progression and metastasis in solid tumors^35-37^; down-miRs: miR-17-5p, miR-34c-5p, miR-203b-5p; up-miRs: miR-135b-5p, miR-155-5p. We utilized three cancer cell lines from two solid tumor types to probe the generality of our data: melanoma (MeWo and 131/4-5B1 (5B1)^38^) and breast cancer (MCF-7). Cells were treated with miRNA mimics for 48h prior to lysis and analysis by Western blot (Supplementary Fig. 3). Given that the “non-targeting control” miRNA mimic was found to alter expression of our sensor in a 3’UTR dependent manner, we were concerned that this would impact CD98hc levels and could not use this as the control for other analysis. Therefore, we tested several miRNAs that had no impact on the 3’UTR CD98hc sensor, to use as an alternate non-targeting control (NTC, Supplementary Fig. 4). Of the three miRNA tested, miR-548ab was chosen as our new NTC because there are no biological processes currently associated with this miRNA^39^. In line with our miRFluR data, down-miRs resulted in a decrease in endogenous CD98hc expression (∼0.5-fold), and up-miRs resulted in an increase in protein levels (Fig. 2 a-d, Supplementary Fig. 5a-b, 6, 1.6-2-fold) in all three cell lines. We next analyzed CD98hc cell-surface expression in MeWo and 5B1 using flow cytometry. Our data again confirmed that up-miRs (miR-135b-5p, miR-155-5p) enhance cell surface expression of CD98hc, while down-miRs (miR-203b-5p) decrease it (Fig. 2e-h). We next analyzed the impact of miR mimics on transcript levels of CD98hc (*SLC3A2*) in MCF-7 and 5B1 (Supplementary Fig. 5c-d, Supplementary Fig. 7). We, and others, have found discordance between the impacts of miRNA on the transcript levels and the observed impacts on protein expression^5,6,40-42^. In keeping with this previous work, we found a mixture of impacts on the transcriptome with little concordance between the transcript and protein levels for either down- or up-miRs. Overall, our data supports our miRFluR analysis, identifying both down- and up-regulators of CD98hc.

**Fig 2.**
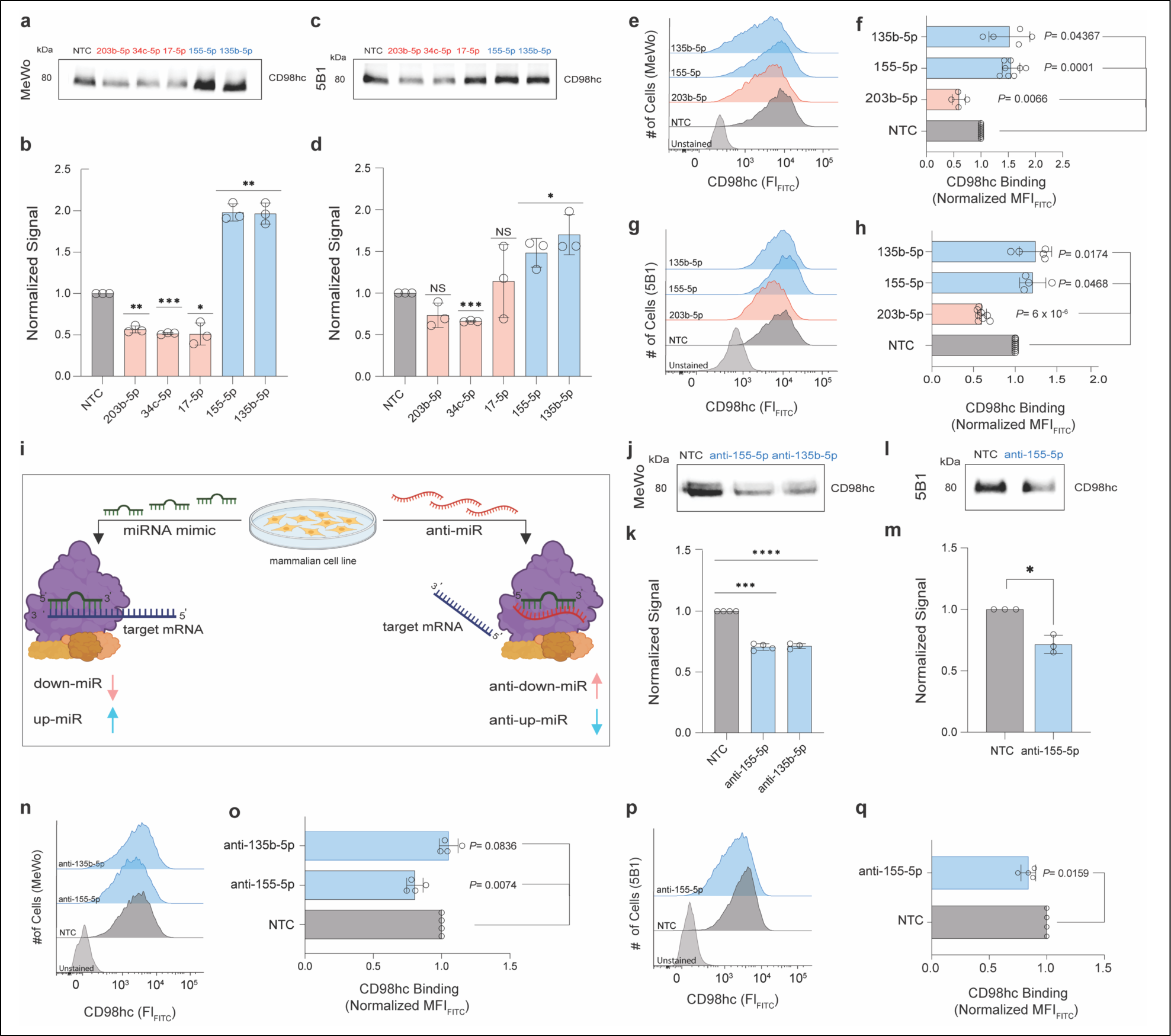
CD98hc miRFluR analysis identifies endogenous regulators of expression. Impact of miRNA mimics on CD98hc expression. MeWo (a, b, e, f) or 5B1 (c, d, g, h) cells were treated with miRNA mimics (down-miRs: miR-203b-5p, -34c-5p, -17-5p, upmiRs: miR-155-5p, -135b-5p) or non-targeting control (NTC: miR-548ab was used for all miR mimic experiments) at 50 nM for 48h and analyzed as indicated. **a, c.** Representative Western blot analysis of CD98hc in MeWo (a) and 5B1 (c). **b, d.** Bar graph of Western blot data for MeWo (b) and 5B1 (d). **e, g**. Representative flow cytometry analysis of CD98hc binding in MeWo (e) and 5B1 (g). **f, h**. Bar graph of flow cytometry data for MeWo (f) and 5B1 (h). Impact of endogenous miRNA on CD98hc expression (i-q). **i**. Anti-miRs sequester the endogenous miRNA by interacting with target in AGO2 complexes, causing loss of protein expression for up-miRs. MeWo (j, k, n, o) or 5B1 (l, m, p, q) cells were treated with anti-miRs (anti-up-miRs: anti-155-5p, anti-135b-5p) or non-targeting control (NTC) at 50 nM for 48h and analyzed as indicated. **j, l**. Representative Western blot analysis of CD98hc in MeWo (j) and 5B1 (l) cells. **k, m**. Bar graph of Western blot data for MeWo (k) and 5B1 (m). **n, p**. Representative flow cytometry analysis of CD98hc binding in MeWo (n) and 5B1 (p). **o, q**. Bar graph of flow cytometry data for MeWo (o) and 5B1 (q). Additional data are shown in Supplementary Fig.s 2-8. All experiments were performed in β biological triplicate. Errors shown are standard deviations. For Western blot analysis, the Wilcoxon *t-*test was used to compare miRs to NTC (ns not significant, * *p < 0.05, ** < 0.01, *** < 0.001*). In flow cytometry analysis, paired *t*-test was used to compare miRs to NTC and *p*-values are indicated on the graph.

To test whether endogenous up-miRs enhanced CD98hc expression, we used miRNA hairpin inhibitors, which displace target mRNA from AGO complexes (anti-miRs, Fig. 2i). These inhibitors should repress expression of proteins whose levels are enhanced by upregulatory miRNA. We focused on up-miRs-155-5p and -135b-5p, both of which are expressed in the melanoma line MeWo^32,43^ (Supplementary Fig. 8). Note, for all anti-miR experiments the anti-miR NTC supplied by the company was used as this is distinct from the mIRIDIAN mimic NTCs. As expected, transfection with anti-up-miRs for miR-155-5p inhibited expression of CD98hc as observed in both Western blot analysis and flow cytometry for MeWo and 5B1 (Fig. 2j-q, Supplementary Fig. 5). However, we observed cell line dependent differences for the anti-miRs of miR-135b-5p, with MeWo showing clear loss of expression and no impact in 5B1. This discrepancy may be due to the expression level of this miRNA in the two cell lines. Overall, our data confirms the validity of our miRFluR assay to identify endogenous miRNA regulating CD98hc.

### miR-155-5p upregulates CD98hc via direct miR: mRNA interactions

The oncogenic roles of CD98hc dovetail with those of miR-155-5p, which is highly expressed in many cancers ^44,45^ ^46,47^. Both are known to be upregulated and functionally important in B-cell lymphoma ^48^, acute myelogenous leukemia^21,49^, and non-small cell lung carcinoma^19,50^. Both are also critical to clonal expansion of B- and T-cells^14-16,51,52^. We anticipated that miR-155-5p would directly bind to the CD98hc 3’UTR to mediate upregulation. To test this, we utilized our miRFluR system (Fig. 3a). In our previous work, the RNAhybrid algorithm was able to accurately predict regulatory binding sites for up-miRs^6^. This algorithm predicts the most stable binding site between miRNA and mRNA^53^. A non-canonical miR-155-5p binding site was identified at position 152 in the CD98hc 3’UTR (Fig. 3b). Using site-directed mutagenesis, we mutated all interacting nucleotides at the potential binding site in our pFmiR-CD98hc sensor and tested the mutant in our assay (Fig. 3c). As expected, we lost miR-155-5p mediated upregulation in the mutant sensor, confirming that upregulation of CD98hc by this miRNA is through direct interactions. In line with our previous work^6^, the upregulatory sites are not AU rich^7,8^, and do not adhere to the seed rules that are often observed in downregulatory sites^3^. Our work suggests that miR-155-5p may be directly involved in the upregulation of CD98hc observed in cancer and immunity.

**Fig. 3.**
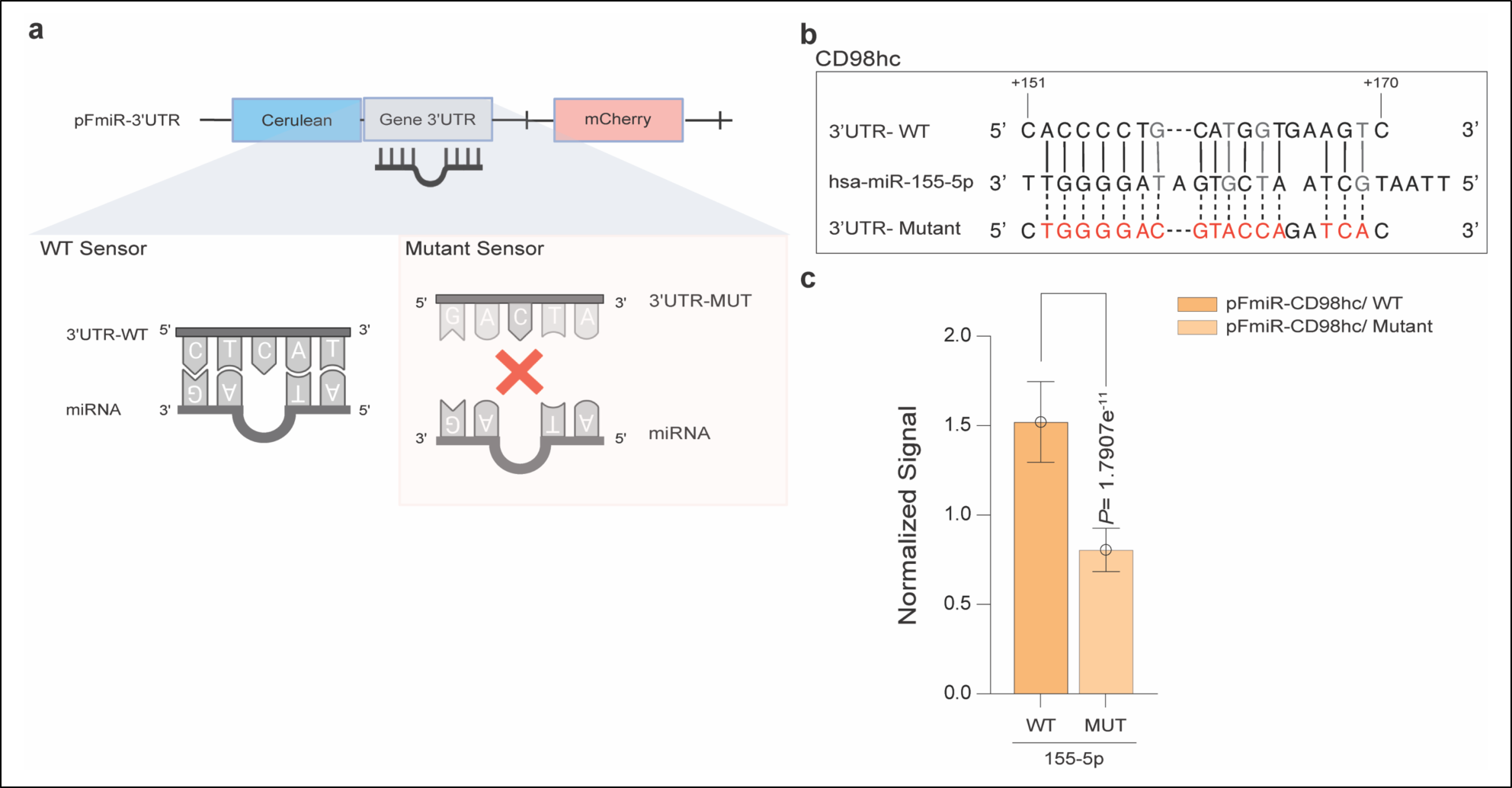
Mutational analysis identifies miR-155-5p binding site on CD98hc 3’UTR. **a.** Comparison of wild-type (WT) and mutant pFmiR-3’UTR interactions with miRNA. **b**. Alignment of miR-155-5p with predicted CD98hc-3′UTR site and corresponding mutant. Mutated residues are shown in red, wobble interactions (G: U) are shown in grey. **c**. Bar graph of data from wildtype (WT) and mutant (MUT) miRFluR sensors. For each sensor, data was normalized over NTC. Experiment was performed in biological duplicate, representative assay with n=8 wells is shown. Standard *t-*test was used for comparison, *p-*value is indicated on graph.

### Mapping miRNA regulation of ST3GAL1&2, mediators of CD98hc stability in melanoma

Co-regulation of critical proteins within a network through coordinated targeting by miRNA has been widely observed for down-miRs^5,40^. There are no existing examples of co-upregulation mediated directly by the miRNA in such networks. In analyzing the up-miRs for CD98hc, we noticed that a significant number of them (15/35, 43%) were associated with melanoma genesis and progression^31^(Supplementary Table 1). CD98hc is highly expressed in metastatic melanoma and is associated with lower survival of melanoma patients^20,32^. In recent work, we identified a role for α-2,3-sialylation, mediated by the sialyltransferases ST3GAL1 and ST3GAL2, in the stabilization of CD98hc in melanoma. Inhibition of ST3GAL1 or ST3GAL2 reduced CD98hc protein stability^11^. Given the impact of ST3GAL1 and ST3GAL2 on CD98hc expression in melanoma, we hypothesized that these genes might be co-upregulated by miRNA. As a first step towards testing this hypothesis, we mapped the miRNA regulation of ST3GAL1 and ST3GAL2 using our miRFluR assay (Figs. 4 and 5).

**Fig. 4.**
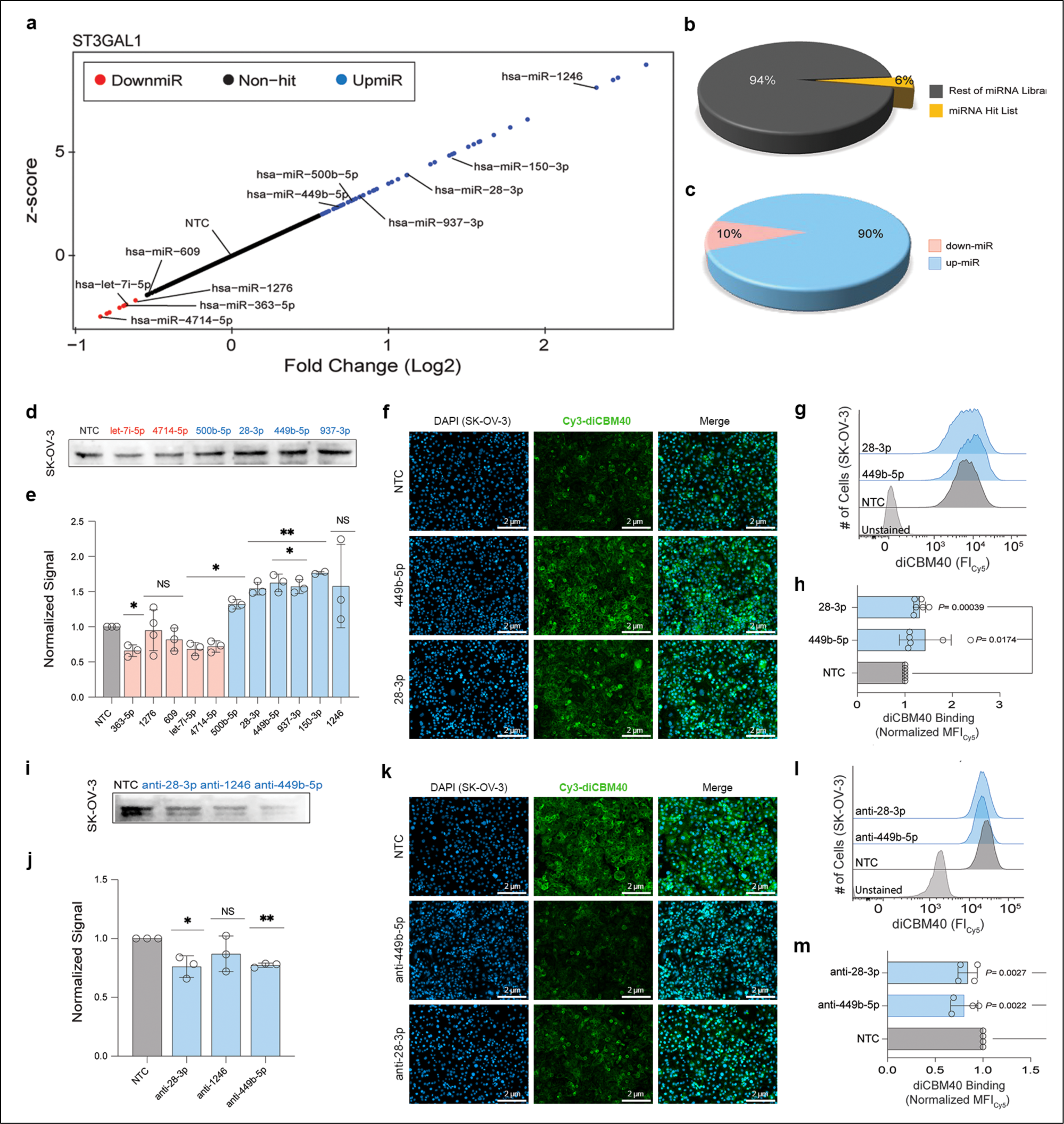
miRNA regulation of ST3GAL1 impacts α-2,3-sialylation. **a.** Landscape of miRNA regulation of ST3GAL1 via the 3’UTR is shown via scatterplot of miRFluR data. miRNA in the 95% confidence interval by z-score are indicated (down-miRs: red, up-miRs: blue) and validated miRNAs are labeled. **b.** Pie chart showing % miRNA hits compared to total library post-QC. **c.** Pie chart indicates % down-miR (10%) vs. up-miR (90%) in ST3GAL1 hit list. **d-h.** Impact of miRNA mimics on ST3GAL1 expression. SK-OV-3 cells were treated with miRNA mimics (down-miRs: -363-5p, -1276, -609, let-7i-5p, -4714-5p, up-miRs: -500b-5p, -28-3p, -449b-5p, -937-3p, - 150-3p, -1246) or non-targeting control (NTC) at 50 nM for 48h and analyzed as indicated. **d**. Representative Western blot analysis of ST3GAL1. **e**. Bar graph of Western blot data. **f.** Fluorescence microscopy for Cy3-labeled diCBM40 staining of SK-OV-3 treated with NTC or up-miRs: -449b-5p and -28-3p. Representative images are shown. **g**. Flow cytometry analysis of Cy-5 labeled diCBM40 binding for SK-OV-3 treated as in f. **h.** Bar graph of flow cytometry data. MFI: mean fluorescence intensity; FI_cy5_: fluorescence intensity. **i-m.** Impact of endogenous miRNA on ST3GAL1 expression. SK-OV-3 cells were treated with anti-miRs (anti-up-miRs: anti-28-3p, anti-1246, anti-449b-5p) or non-targeting control (NTC) at 50 nM for 48h and analyzed as indicated. **i**. Representative Western blot analysis of ST3GAL1. **j**. Bar graph of Western blot data. **k.** Fluorescence microscopy analysis for diCBM40 staining. **l.** Representative flow cytometry analysis of diCBM40 binding. **m.** Bar graph of flow cytometry data. Additional data are shown in Supplementary Figs 9-12. All experiments were performed in β biological triplicate. Errors shown are standard deviations. For Western blot analysis, the Wilcoxon *t-*test was used to compare miRs to NTC (ns not significant, *\* p < 0.05, ** < 0.01, *** < 0.001*). For flow cytometry analysis, the paired *t*-test was used to compare miRs to NTC and *p*-values are indicated on the graph.

**Fig. 5.**
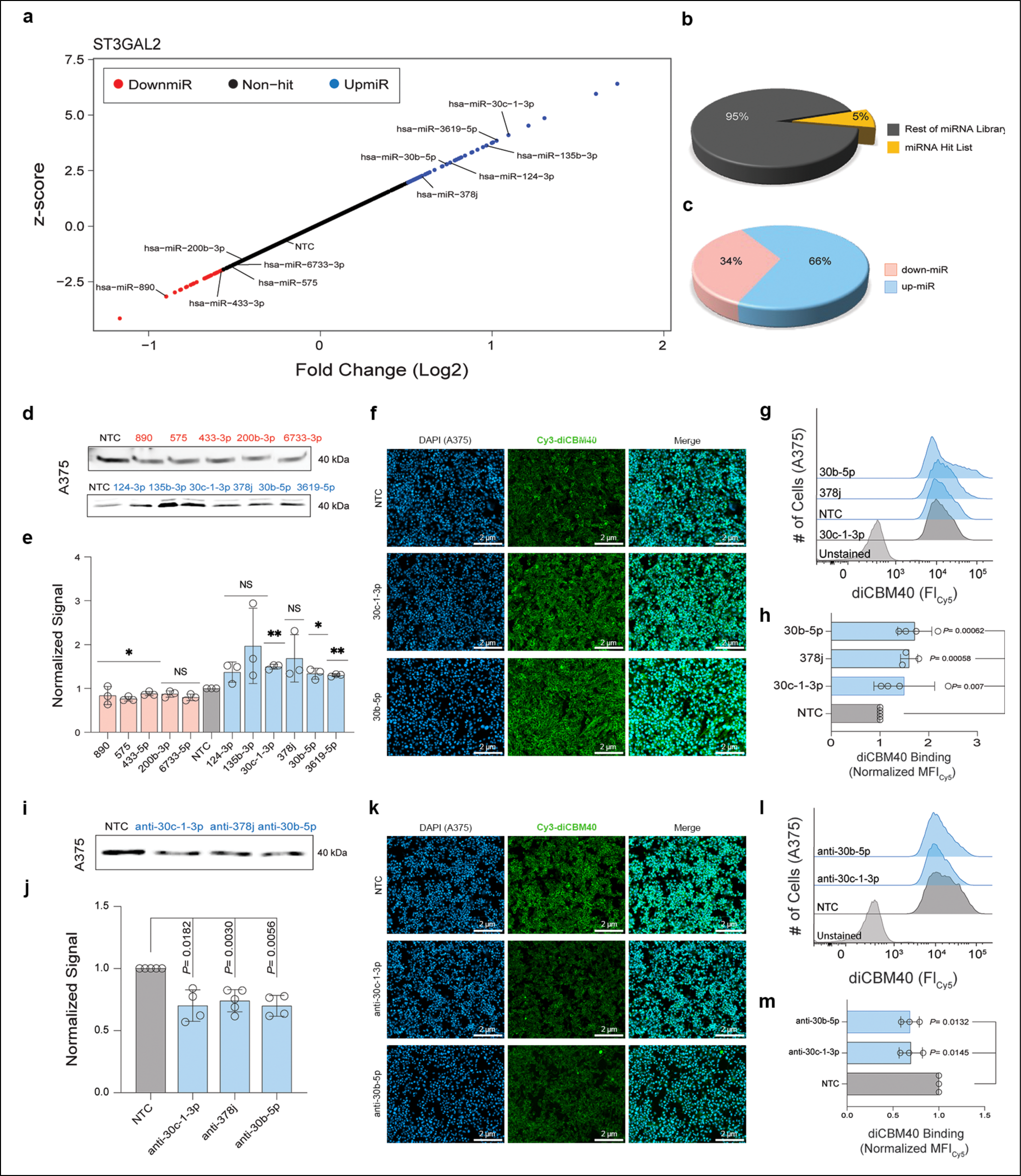
miRNA regulation of ST3GAL2 impacts α-2,3-sialylation. **a.** Landscape of miRNA regulation of ST3GAL2 via the 3’UTR is shown via scatterplot of miRFluR data. miRNA in the 95% confidence interval by z-score are indicated (down-miRs: red, up-miRs: blue) and validated miRNAs are labeled. **b.** Pie chart showing % miRNA hits compared to total library post-QC. **c.** Pie chart indicates % down-miR (34%) vs. up-miR (66%) in ST3GAL2 hit list. **d-h.** Impact of miRNA mimics on ST3GAL2 expression. A375 cells were treated with miRNA mimics (down-miRs: -890, -575, -433-5p, -200b-3p, -6733-5p, upmiRs: -124-3p, -135b-3p, -30c-1-3p, -378j, - 30b-5p, -3619-5p) or non-targeting control (NTC) at 50 nM for 48h and analyzed as indicated. **d**. Representative Western blot analysis of ST3GAL2. **e**. Bar graph of Western blot data. **f.** Fluorescence microscopy analysis for Cy3-diCBM40 staining. **g**. Representative flow cytometry analysis of Cy5-diCBM40 binding. **h**. Bar graph of flow cytometry data. **i-m.** Impact of endogenous miRNA on ST3GAL2 expression. A375 cells were treated with anti-miRs (anti-up-miRs: anti-30c-1-3p, anti-378j, anti-30b-5p) or non-targeting control (NTC) at 50 nM for 48h and analyzed as indicated. **i**. Representative Western blot analysis of ST3GAL2. **j**. Bar graph of Western blot data. **k.** Fluorescence microscopy analysis for of Cy3-diCBM40 staining. **l.** Representative flow cytometry analysis of Cy5-diCBM40 binding. **m.** Bar graph of flow cytometry data. Additional data are shown in Supplementary Figs 13-16. All experiments were performed in ≥ biological triplicate. Errors shown are standard deviations. For Western blot analysis the Wilcoxon *t-*test was used to compare miRs to NTC (ns not significant, * *p < 0.05, ** < 0.01, *** < 0.001*). For flow cytometry analysis, the paired *t*-test was used to compare miRs to NTC and *p*-values are indicated on the graph.

### High-throughput analysis shows ST3GAL1 is predominantly up-regulated by miRNA

ST3GAL1, one of six glycosyltransferases that transfer sialic acid to the 3-position of terminal galactose creating the α−2,3-sialyl epitope, predominantly sialylates *O*-glycans such as those found on mucins^23^. It impacts both growth and metastasis in melanoma and alters T-cell migration^11,27,28^. ST3GAL1 upregulation also plays a significant role in the development of malignant epithelial ovarian cancer^26^ and colon cancer^54,55^. We cloned the most prevalent 3’UTR of ST3GAL1 (∼5 kb) into our pFmiR sensor (pFmiR-ST3GAL1, Supplementary Fig. 9) for analysis in our miRFluR assay. Of the 1807 miRNAs that passed QC, we identified 7 down-miRs and 60 up-miRs within the 95% confidence interval (Fig. 4a-c, Supplementary Fig. 10, Dataset 2). Our NTC (miR-548ab) was again in the middle of our dataset and did not impact ST3GAL1 levels. Interestingly, the majority (90%) of miRNA hits were upregulatory, which is similar to our observations for ST6GAL1^6^. We chose miRNA that were biologically significant and spread out within the 95% confidence interval for validation. We focused on 5 down-miRs (miR-363-5p, miR-1276, miR-609, miR-4714-5p, let-7i-5p) and 6 up-miRs (miR-937-3p, miR-500b-5p, miR-28-3p, miR-449b-5p, miR-1246, miR-150-3p), the majority of which were associated with various forms of cancer biology^56-59^. Because ST3GAL1 is highly expressed in ovarian and colon cancers, we initially tested our miRNA in the ovarian cancer cell line (SK-OV-3) and the colon cancer cell line (HT-29) for their impact on ST3GAL1 (Fig. 4, Supplementary Figs. 11-12). As expected, Western blot analysis of ST3GAL1 in cells treated with miRNA mimics followed the regulation observed in our miRFluR assay (Fig. 4d-e, Supplementary Fig. 11a-b). In keeping with previous work, transcript levels did not directly follow protein levels (Supplementary Fig. 11c-d)^5,6,40-42^. We next tested the potential impact of select up-miRs (miRs -28-3p, -449b-5p) on *α*-2,3-sialylation in SK-OV-3 using the Cy3-conjugated lectin diCBM40^60^. In line with our expectations, we observe upregulation of *α*-2,3-sialic acid by both fluorescence microscopy and flow cytometry (Figs. 4f-h, Supplementary Fig. 11e). To confirm that ST3GAL1 is regulated by the corresponding endogenous miRNA, we transfected SK-OV-3 cells with anti-up-miRs -449b-5p, -28-3p and -1246. We observed a loss of both ST3GAL1 protein expression and the corresponding *α*-2,3-sialic acid (Fig. 4i-m, Supplementary Fig. 11f). Our data again confirms the validity of our miRFluR assay and provides strong evidence for endogenous upregulation of ST3GAL1.

### Mapping the miRNA regulatory landscape of ST3GAL2

ST3GAL2, another member of the *α*-2,3-sialyltransferase family, is thought to modify both *O*-glycans and glycolipids^22^. Overexpression of ST3GAL2 may help drive tumorigenesis in several cancers^25,29^. We mapped the miRNA regulation of the most prevalent 3’UTR of ST3GAL2 using our miRFluR assay (pFmiR-ST3GAL2, Supplementary Fig. 13). After data processing, we identified 28 down-miRs and 56 up-miRs within the 95% confidence interval (Fig. 5 a-c, Supplementary Fig. 14, Dataset 3). As before, our NTC (miR-548ab) was again in the middle of our dataset and did not impact ST3GAL2 levels. We again chose a subset of miRNA (down-miRs: miR-890, miR-433-3p, miR-575, miR-6733-3p, miR-200b-3p; up-miRs: miR-124-3p, miR-3619-5p, miR-135b-3p, miR-30c-1-3p, miR-378j, miR-30b-5p) based on the criteria outlined above for validation^31,61^. miRNA mimics were transfected into A375 (melanoma, Fig. 5d-e, Supplementary Figs. 15, 16a-f) or A549 (lung carcinoma, Supplementary Figs. 15a-b, 16g-j) and analyzed by Western blot analysis after 48 hours. The resulting impacts on ST3GAL2 protein expression were in line with our miRFluR assay. In contrast, the effect of miRNA mimics on the transcript of *st3gal2* was variable, as previously noted for other genes (Supplementary Fig. 15c-d). We next tested the effect of select up-miRs (miRs -30b-5p, -30c-1-3p, -378j) on α-2,3-sialic acid levels in A375 using fluorescence microscopy and flow cytometry and observed the anticipated increase in sialylation (Fig. 5f-h, Supplementary Fig. 15e-f). We confirmed the impact of endogenous upregulatory miRNA using anti-miRs (anti-up-miRs -30b-5p, -30c-1-3p, -378j) and observed the expected loss of both protein and glycan expression (Fig. 5i-m). Our data again showcases the power of our miRFluR assay to explore the miRNA regulatory landscape.

### miRNA co-upregulate CD98hc and α-2,3-sialylation in Melanoma

Given the importance of ST3GAL1 and ST3GAL2 on CD98hc stability in melanoma^11^, we examined whether they might be co-regulated by miRNA. Comparison of all three miRNA datasets did not identify miRNA that regulated all three proteins. We also did not observe miRNA that have opposing effects (i.e., up-miR in one, down-miR in another) between any of the three. However, CD98hc and ST3GAL1 shared 6 up-miRs in common (miR-765, <u>-488-5p</u>, -<u>449b-5p</u>, -500b-5p, <u>-28-3p</u>, <u>miR-1246</u>), <u>4</u> of which were upregulated in melanoma during either the transformation of benign nevi^62^ or progression (underlined)^32^. In addition, four up-miRs were found to co-regulate CD98hc and ST3GAL2 (miR-378j, <u>-135b-3p</u>, <u>-30b-5p</u>, <u>-30c-1-3p</u>), <u>3</u> of which were associated with melanoma^31,32,61,62^ (Fig. 6). To address whether these miRs might co-regulate CD98hc and sialylation by ST3GAL1 or ST3GAL2, we tested the impact of a select subset of these co-regulatory miRs (ST3GAL1: miR-1246, -28-3p, -500b-3p, -449b-5p; ST3GAL2: miR-378j, -135b-3p, -30b-5p, -30c-1-3p) on their expression in the melanoma cell lines MeWo and 5B1 and the breast cancer line MCF-7 (Fig. 7, Supplementary Figs. 17-21). In line with our miRFluR results, transfection of co-upmiRs upregulated both CD98hc (orange) and either ST3GAL1 (purple) or ST3GAL2 (green) in all three cell lines as seen by Western blot analysis (Fig. 7a-d, Supplementary Figs. 18a-d, 20). Flow cytometry analysis of CD98hc confirmed Western blot results (Fig. 7e-f, Supplementary Figs. 18e-f). We also observed an increase in α-2-3-sialylation by flow cytometry, in line with upregulation of ST3GAL1 or ST3GAL2 expression by the up-miRs (Fig. 7g-h, Supplementary Figs. 18g-h). To confirm that endogenous miRNA could co-regulate these proteins, we utilized the corresponding anti-miRs to the co-upregulators (Fig. 7i-p, Supplementary Fig. 18i-p). All miRNA tested were known to be expressed in either melanoma or the MeWo cell line^32,43^ (Supplementary Fig. 8). As anticipated, Western blot analysis clearly shows loss of expression for co-upregulated proteins upon inhibition of the endogenous miRNA. This data was supported by flow cytometry analysis of CD98hc. We also observed the loss of α-2,3 sialic acid, concordant with the loss of ST3GAL1 or ST3GAL2 expression upon up-miR inhibition, although this was only statistically significant in the 5B1 cell line (Supplementary Fig. 18p). Overall, our data strongly supports the idea that miRNA regulatory networks include co-upregulation of critical proteins.

**Fig. 6.**
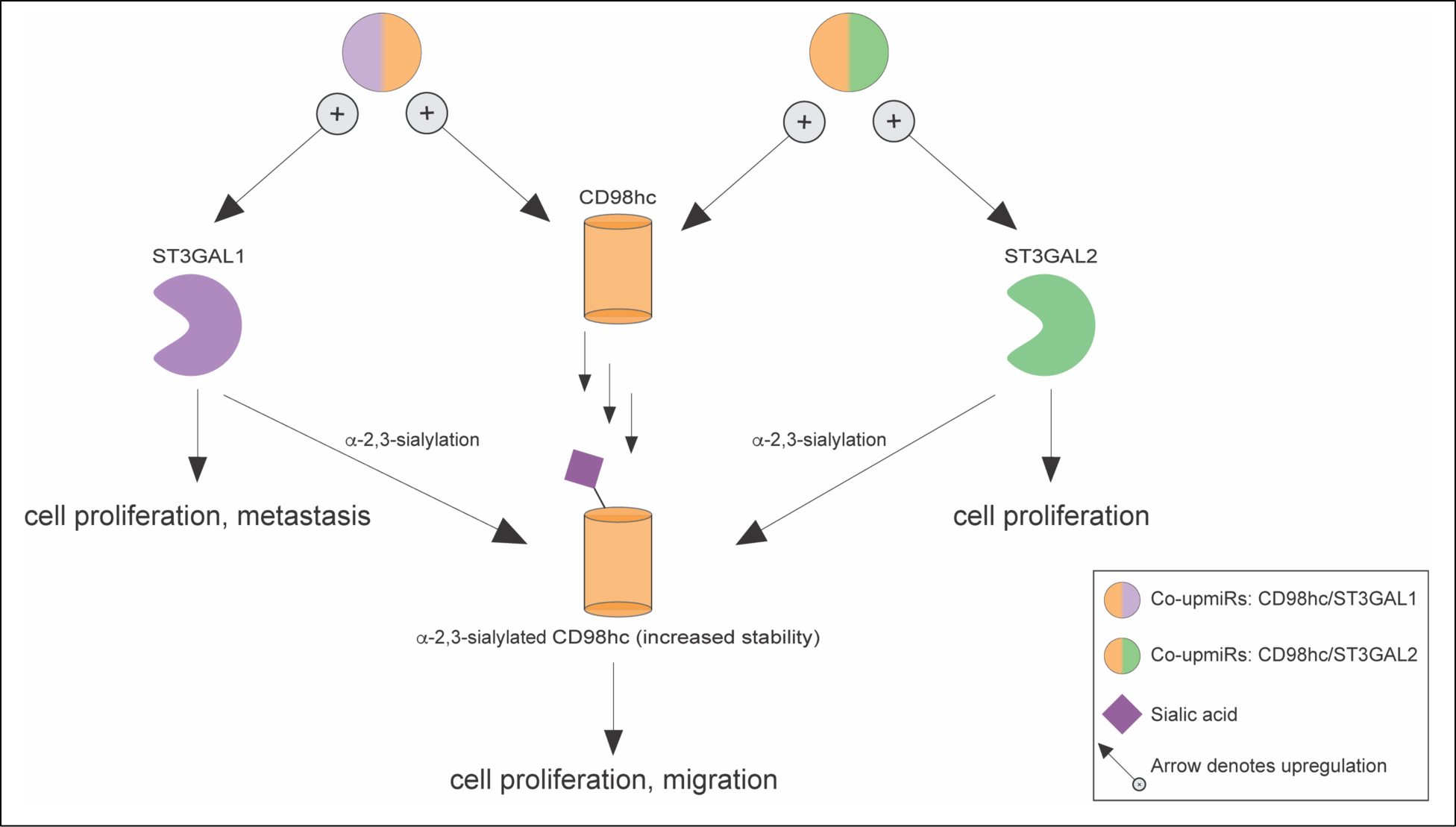
Upregulatory miRNAs co-regulate CD98hc/ST3GAL1 or CD98hc/ST3GAL2. Sialylation is required for the stability of CD98hc in melanoma where ST3GAL1, ST3GAL2 and CD98hc are all essential genes.

**Fig. 7.**
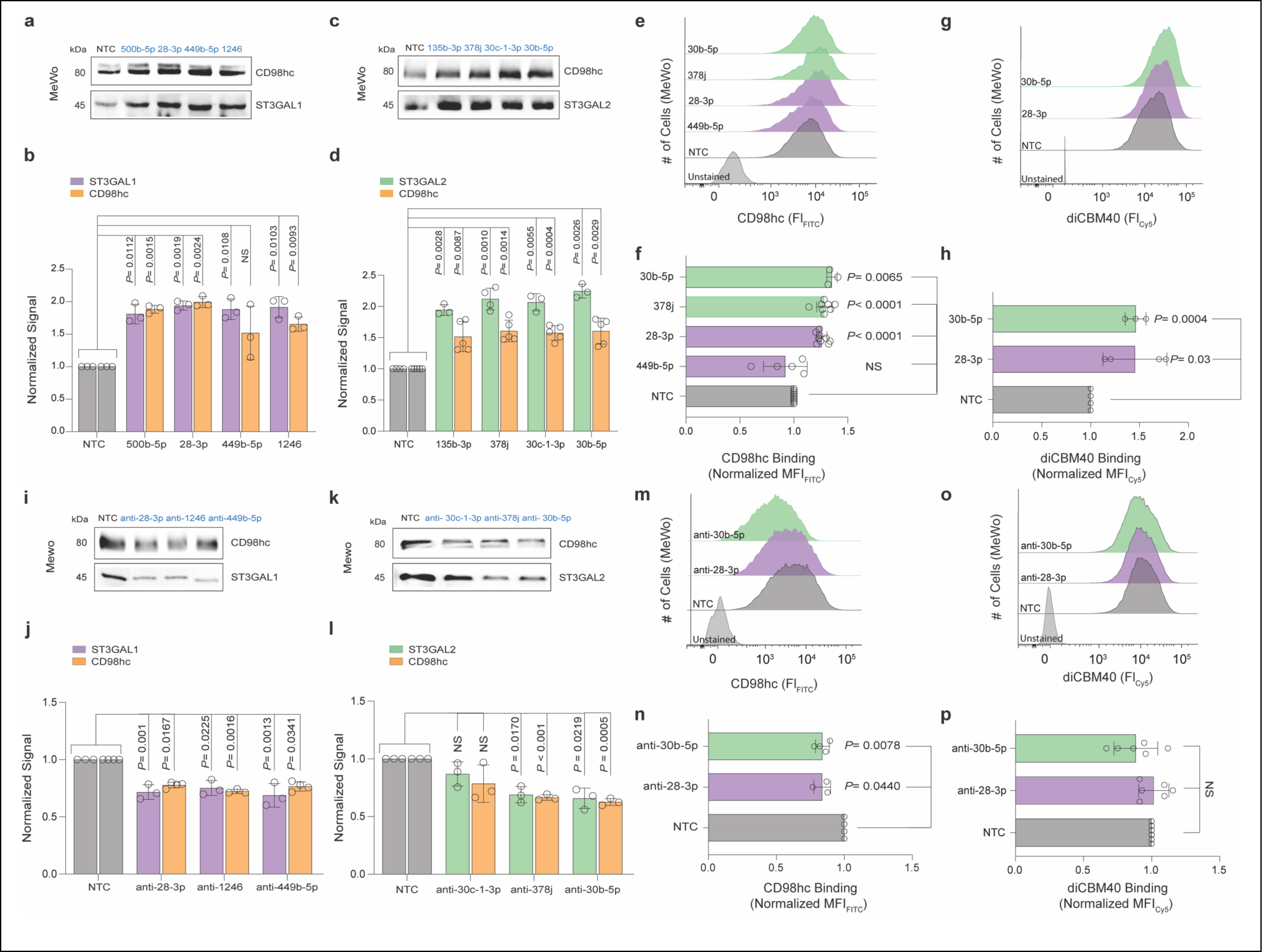
Co-upregulation of CD98hc and either ST3GAL1 or ST3GAL2 is observed in melanoma. **a-d.** Impact of co-up-miR mimics on either CD98hc (orange) & ST3GAL1 (purple, co-up-miRs: -500b-5p, -28-3p, -449b-5p, - 1246) or CD98hc (orange) & ST3GAL2 expression (green, co-up-miRs: -135b-3p, -378j, -30c-1-3p, -30b-5p) in MeWo cells treated with corresponding co-up-miRs or non-targeting control (NTC) at 50 nM for 48h and analyzed as indicated. **a**. Representative Western blot analysis of CD98hc and ST3GAL1. **b**. Bar graph of corresponding Western blot data. **c**. Representative Western blot analysis of CD98hc and ST3GAL2. **d**. Bar graph of corresponding Western blot data. **e**. Representative flow cytometry analysis of CD98hc binding. **f**. Bar graph of flow cytometry data. **g**. Representative flow cytometry analysis of Cy5-diCBM40 binding. **h**. Bar graph of flow cytometry data. **i-o.** Impact of endogenous co-up-miRs on either CD98hc (orange) & ST3GAL1 (purple) or CD98hc (orange) & ST3GAL2 (green) expression. MeWo cells were treated with anti-miRs (anti-up-miRs: anti-28-3p, anti-1246, anti-449b-5p; anti-30c-1-3p, anti-378j, anti-30b-5p) or non-targeting control (NTC) at 50 nM for 48h and analyzed as indicated. **i**. Representative Western blot analysis of CD98hc and ST3GAL1. **j**. Bar graph of Western blot data. **k**. Representative Western blot analysis of CD98hc and ST3GAL2. **l**. Bar graph of Western blot data. **m.** Representative flow cytometry analysis of CD98hc binding. **n.** Bar graph of flow cytometry data for MeWo. **o.** Representative flow cytometry analysis of Cy5-diCBM40 binding. **p.** Bar graph of flow cytometry data. Additional data are shown in Supplementary Figs 17-21. All experiments were performed in ≥ biological triplicate. Errors shown are standard deviations. For Western blot analysis, the Wilcoxon *t-*test was used to compare miRs to NTC (ns not significant, *\* p < 0.05, ** < 0.01, *** < 0.001*). For flow cytometry analysis, paired *t*-test was used to compare miRs to NTC and *p*-values are indicated on the graph.

### Co-regulation of CD98hc and α-2,3-sialic acid is via direct miRNA:3’UTR interactions

The impact of miRNA on protein expression can be via direct or indirect effects. In our previous works, we established that our miRFluR assay identifies miRNA regulation via direct interactions between the miRNA and the 3’UTR of the transcript of interest^5,6^. To verify that miRNA co-upregulation of CD98hc and either ST3GAL1 or ST3GAL2 were via direct miRNA:mRNA interactions, we identified the upregulatory sites for two miRNAs: miR-1246 (CD98hc/ST3GAL1) and miR-30b-5p (CD98hc/ST3GAL2) using RNAhybrid^53^. These miRNAs are highly expressed in melanoma and are associated with melanoma progression^31,61^. We then mutated these sites in our pFmiR sensors and tested their impact on miRNA-mediated upregulation (Fig. 8). In line with our earlier work, mutation of the RNAhybrid predicted sites abrogated upregulation. We also identified and validated the site for miR-28-3p (a co-up-miR for ST3GAL1/CD98hc) for CD98hc (Supplementary Fig. 22). None of the sites identified had canonical seed binding, nor were they AU rich. Taken together, our data strongly supports an alternative binding motif for upregulatory miRNA:mRNA interactions that has yet to be fully determined.

**Fig. 8.**
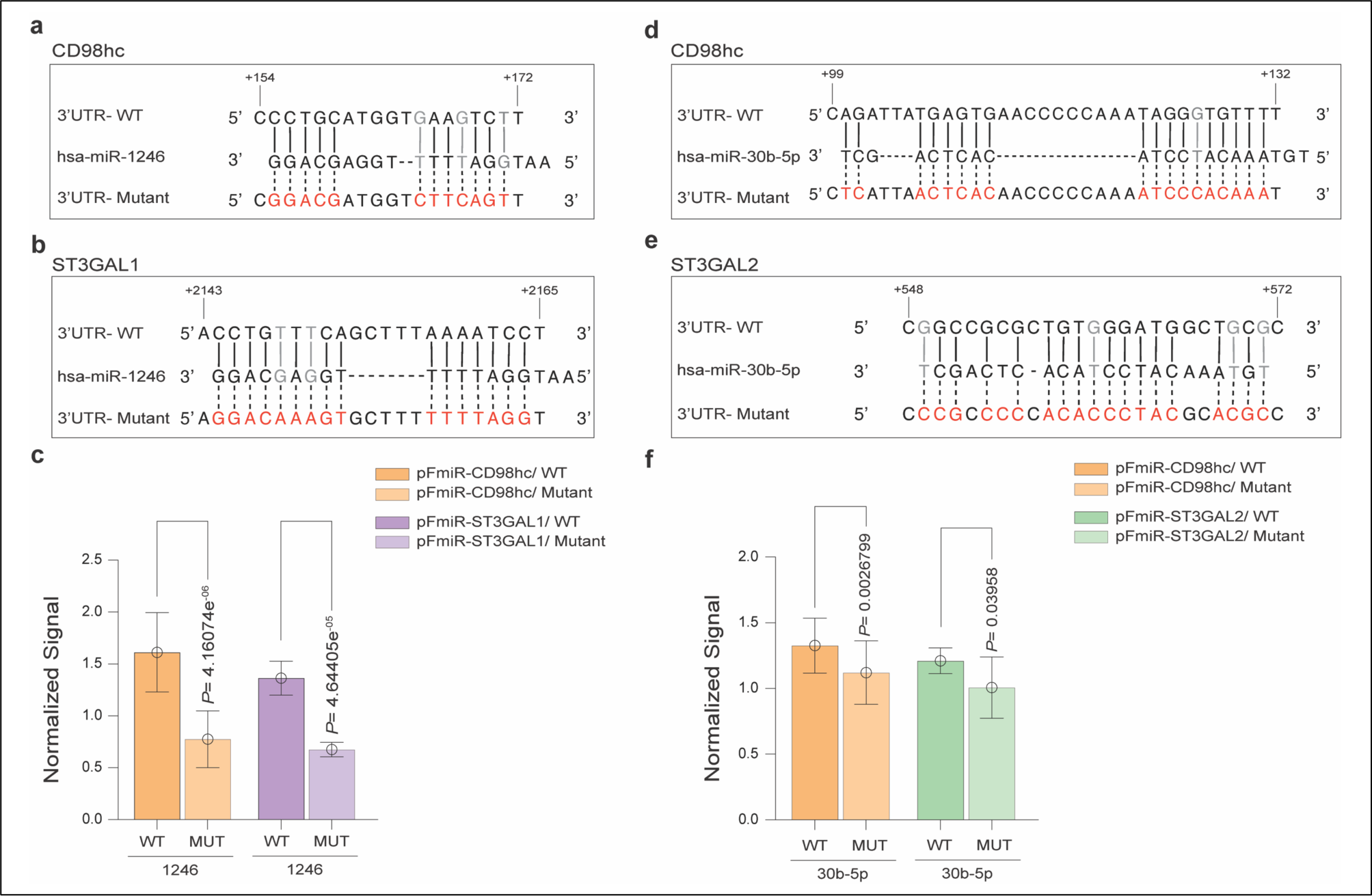
Co-upregulation by miRNAs requires direct interactions with 3′UTRs of CD98, ST3GAL1 and ST3GAL2. **a, b**. Alignment of miR-1246 with predicted CD98hc-3′UTR (a) or ST3GAL1-3’UTR (b) sites and their corresponding mutants. Mutated residues are shown in red. Wobble interactions are shown in grey. **c**. Bar graph of data from miRFluR sensors. For each sensor, data was normalized over NTC. **d, e**. Alignment of miR-30b-5p with predicted CD98hc-3′UTR (d) or ST3GAL2-3’UTR (e) sites and their corresponding mutants. Mutated residues are shown in red. Wobble interactions are shown in grey. **f**. Bar graph of data from miRFluR sensors. For each sensor, data was normalized over NTC. Propagated error for the sensor data is shown. A standard *t*-test of the NTC normalized data was used for comparisons. All experiments were performed in biological triplicate, a representative experiment with n = 10 wells is shown.

## Discussion

The canonical view of miRNA, which can have >100 targets each, is that they downregulate the expression of proteins that synergistically control specific cellular pathways and phenotypes. In recent work, our laboratory discovered that miRNA can upregulate protein expression through direct binding to the 3’UTR in dividing cells^6^. This contradicts the dominant view of miRNA, providing a mechanism for bidirectional tuning of protein expression. Herein, we provide further evidence for upregulation and expand upon its roles in miRNA-mediated tuning of protein networks.

In mapping the miRNA landscape for both the sialyltransferases (ST3GAL1, ST3GAL2) and their glycosylation target (CD98hc), we identify up- and downregulatory miRNAs for all three genes (up-miRs:151 total interactions, down-miRs: 65 total interactions). As expected, biological validation of up- and down-miRs using both miRs and anti-miRs followed the data from our miRFluR assay. Our previous work focused on identifying regulators of glycosyltransferases, which are often medium to low abundance proteins^5,6^. In contrast, CD98hc (*SLC3A2*) is a component of a neutral amino acid transporter found at the cell surface^12,63^. It is highly abundant in cancer and in proliferating B- and T-cells, where it plays an important role in clonal expansion^14-16^. Of note, one of the strongest hits observed in our assay for CD98hc was miR-155-5p, a highly abundant miRNA which itself is critical in both cancer and clonal expansion of B- and T-cells^34,51,52^. The concordance of roles for miR-155-5p and CD98hc strongly suggests that CD98hc is a primary target of miR-155-5p in these biological systems.

The majority of miRNA regulation is currently thought to occur at the 3’UTR of transcripts, and our miRFluR assay is focused on such regulation. The length of the 3’UTR has increased as organisms have become more complex and required enhanced regulation of protein expression. The median 3’UTR length in humans is ∼1,200 nucleotides^64,65^. Similar to other genes analyzed using our miRFluR assay to date^5,6^, the 3’UTRs of ST3GAL1 and ST3GAL2 are both > 2 kb. However, the 3’UTR of CD98hc is notably short, at only 190 nt. Upregulatory miRNAs were seen for all three genes, indicating that upregulation does not require a long 3’UTR. The proportion of up- to down-miRs for these genes was target dependent, with ST3GAL1 biased towards up-miRs (90%, Fig. 4c) and CD98hc almost evenly divided between up- and down-regulators (54:46, up: down, Fig. 1e). Consistent with these differences, in our previous work, we observed transcripts biased toward down-regulatory miR interactions (ST6GAL2) and those biased towards upregulation (ST6GAL1)^6^. This hints that structure of the 3’UTR may influence the consequence of miRNA interactions, changing the predominant mode of regulation.

The local binding between miRNA and their cognate mRNA in downregulatory complexes is highly enriched in seed sequences, in which positions 2-8 at the 5’-region of the miRNA are base paired to the mRNA^66^. Of the six upregulatory binding sites identified in this work (Figs. 3, 8, Supplementary Fig. 22), none follow canonical seed rules. Our previous work identified 3 upregulatory binding sites, of which only one had a canonical seed^6^. We, and others, found that upregulation by miRNA in both dividing ^6^ and quiescent ^7^ cells requires Argonaute 2 (AGO2), but is inhibited by TNRC6A (GW182), an essential component of downregulatory AGO2 complexes^9,10^. In a re-evaluation of miRNA:mRNA target base-pairing within AGO complexes, Helwak, *et. al.,* found that non-canonical binding was highly prevalent and that there were distinct non-canonical base-pairing patterns^67^. The same miRNA were found to engage in both seed and non-canonically patterned binding. They posited that differential binding patterns of miRNA: mRNA complexes might alter AGO2 structures, leading to different complexes and activities. The data to date supports the hypothesis that the interaction rules between a single miRNA and its cognate transcript are different in up- and downregulation, potentially recruiting distinct components into AGO2 complexes that define the translational fate of the mRNA^6,7^.

Within the cell, miRNA cooperatively tune the expression of proteins that act in concert within a biological pathway^4,68,69^. This role has been well established for downregulatory miRNA interactions. Co-regulation of proteins in a network by up-miRs has yet to be reported. In recent work, we identified ST3GAL1 and ST3GAL2 as essential genes in melanoma. Loss of these genes caused apoptosis and enhanced proteasomal degradation of CD98hc, itself an essential gene^11^. In strong concordance with the biological relationship between CD98hc and ST3GAL1/2 in melanoma, we observed co-upregulators for CD98hc with either ST3GAL1 or ST3GAL2 (Fig. 6). We do not observe any overlap between the miRFluR hits for ST3GAL1 and ST3GAL2, nor do we observe either co-down-miRs or opposing miRNA regulation between the three genes. Of the ten co-upregulators identified, six are overexpressed during melanoma transformation and progression (CD98hc/ST3GAL1: miRs -28-3p, -1246, -449b-5p; CD98hc/ST3GAL2: miRs -135b-3p, -30b-5p, -30c-1-3p)^31,32,61^. This is consistent with the increased protein expression of CD98hc, ST3GAL1 and ST3GAL2 observed in melanoma patients^20,27^. Co-upregulation makes biological sense as α-2,3-sialylation by these enzymes is required for CD98hc protein stability in melanoma, which is most likely mediated by protein glycosylation, a known regulator of trafficking and stability^11,70^.

Several of the miRNA acting as co-upregulators have other known targets in melanoma biology. Proliferation and viability of melanoma cells were enhanced by the expression of miR-1246, which was found to directly target and downregulate the tumor suppressor FOXA2^61^. In earlier work from our laboratory, miR-30b-5p was identified as overexpressed in melanoma metastasis and was found to inhibit the expression of GALNT7, a GalNAc transferase, increasing the invasive capacity of melanoma cells *in vitro* and the metastatic potential *in vivo*^31^. Taken together with our data, this provides strong evidence for bidirectional tuning by the same miRNA within regulatory networks, with upregulation vs. downregulation determined by the miRNA: mRNA pair. Our work positions upregulation as a central part of miRNA regulatory networks and argues for a re-examination of the functions of miRNA.

## Method

### Cloning

CD98hc, ST3GAL1 and ST3GAL2 3’UTRs were amplified via PCR from genomic DNA (gDNA, 10 μg, from MCF-7 cell line: CD98hc) or gene synthesis (GenScript Gene Synthesis company: ST3GAL1 and ST3GAL2) using the primers shown in Supplementary Table 1. The amplicons were cleaned up using a Monarch^®^ PCR & DNA cleanup kit (catalog #: T1030S, NEB). The 3’UTR fragments were then cloned into our pFmiR-empty backbone vector ^5^, downstream of Cerulean with the following restriction sites (for CD98hc & ST3GAL2: NheI and BamHI; for ST3GAL1: AgeI and BsiWI) using standard ligation protocols (NEB). Plasmids were verified by Sanger sequencing (Molecular Biology Services Unit, University of Alberta). Large-scale endotoxin free DNA preparations were made for sequence-verified constructs (pFmiR-ST3GAL1 and pFmiR-ST3GAL2) using Endotoxin-free plasmid DNA purification (Takara Bio USA, Inc., catalog #: 740548). Plasmid maps for pFmiRs -CD98hc, -ST3GAL1 and -ST3GAL2 and their corresponding 3’UTR sequences can be found in Supplementary Figs. 1, 9 and 13.

### Cell Lines

Cell lines HEK-293T, SK-OV-3, A549, and MCF-7 were purchased directly from the American Type Culture Collection (ATCC), HT-29 was a gift from lab of Kristi Baker, MeWo, 131/4-5B1, and A375 were gifts from the Hernando Lab. Cells were cultured in media as follows: HT-29 & HEK-293T:Dulbecco’s Modified Eagle Medium (DMEM), 10% FBS; A549: FK-12, 10%, FBS; A375 & MeWo: DMEM* (catalog # 30-2002), 10% FBS; 131/4-5B1: RPMI-1640, 10% FBS); SK-OV-3: McCoy’s 5a Medium Modified (catalog # 30-2007), 10% FBS; MCF-7: DMEM/F12 (catalog #: 11320033), 10% FBS, under standard conditions (5% CO_2_, 37 °C).

### miRFluR High-throughput Assay

The Human miRIDIAN miRNA mimic Library 21.0 (Dharmacon) was resuspended in ultrapure nuclease-free water (REF. #: 10977-015, Invitrogen) and aliquoted into black 384-well, clear optical bottom tissue-culture treated plates (Nunc). Each plate contained three replicate wells of each miRNA in that plate (2 pmol/well). In addition, each plate contained a minimum of 6 wells containing miRIDIAN non-targeting controls. To each well was added target pFmiR plasmid (pFmiR-CD98hc: 30 ng, pFmiR-ST3GAL1: 30 ng, or pFmiR-ST3GAL2: 20 ng) in 5 µl Opti-MEM (Gibco) and 5 μL of transfection solution (0.1 µL Lipofectamine™ 2000 (catalog #: 11668500, Life Technologies) diluted to 5 µl total volume with Opti-MEM (Gibco), premixed 5 min at room temperature). The mixture was allowed to incubate at room temperature in the plate for 20 min. HEK293T cells (25 µl per well, 400 cells/ µl in phenol red free DMEM supplemented with 10 % FBS and Pen/Strep) were then added to the plate. Plates were incubated at 37°C, 5% CO_2_. After 48 hours, the fluorescence signals of Cerulean (excitation: 433 nm; emission: 475 nm) and mCherry (excitation: 587 nm; emission: 610 nm) were measured using the clear bottom read option (SYNERGY H1, BioTek, Gen 5 software, version 3.08.01).

### Data Analysis

We calculated the ratio of Cerulean fluorescence over mCherry fluorescence (Cer/mCh) for each well in each plate. For each miRNA, triplicate values of the ratios were averaged, and the standard deviation (S.D.) obtained. We calculated % error of measurement for each miRNA (100 × S.D./mean). As a quality control measurement (QC), we removed any plates or miRNAs that had high errors in the measurement (median error ±2 S.D. across all plates) and/or a high median error of measurement for the plate (>15%). After QC we obtained data for 954 miRNAs for CD98hc, 1,807 miRNAs for ST3GAL1, and 1,542 miRNAs for ST3GAL2 out of 2601 total miRNAs screened for each target 3’UTR. We checked whether the miRIDIAN non-targeting controls were at the median and found that they were skewed. Therefore, the Cer/mCh ratio for each miRNA was normalized to the median of Cer/mCh ratios within that plate without exclusions of any data in the plate and error was propagated. Data from all plates were then combined and log-transformed z-scores calculated. A z-score of ±1.965, corresponding to a *two*-tailed *p*-value of 0.05, was used as a threshold for significance. Post-analysis we identified 65 miRNA hits for CD98hc, 67 miRNA hits for ST3GAL1, and 84 for ST3GAL2 within 95% confidence interval (see Fig.s 1c, 4a, 5a, Supplementary Fig.s 2, 10, 14., and Datasets 1-3 (.xls sheet)).

### Western Blots: CD98hc, ST3GAL1 and ST3GAL2

All primary antibodies used in Western blot analysis were validated using pooled siRNA (Dharmacon) and standard conditions (CD98: Supplementary Fig. 3; ST3GAL1: Supplementary Fig. 12, ST3GAL2-Reference 11). Western blot analysis was conducted for CD98hc in MeWo, 5B1 and MCF-7 cell lines (down-miRs: miRs -17-5p, -34c-5p, - 203b-5p; up-miRs: miRs -135b-5p, -155-5p). For ST3GAL1, Western blot analysis was done in SK-OV-3 and HT-29 cell lines (down-miRs: miRs -363-5p, -1276, -609, -4714-5p, let-7i-5p; up-miRs: miRs -937-3p, -500b-5p, -28-3p, -449b-5p, -1246, -150-3p). For ST3GAL2, the Western blot analysis was performed in A375 and A549 cell lines (down-miRs: miRs -890, -433-3p, -575, -6733-3p, -200b-3p; up-miRs: miRs -124-3p, -3619-5p, -135b-3p, -30c-1-3p, -378j, -30b-5p). For co-up-miRs (CD98hc/ST3GAL1: miRs -1246, - 28-3p, -449b-5p, -500b-5p; CD98hc/ST3GAL2: miRs -30b-5p, -30c-1-3p, -378j, -135b-3p), Western blot analysis was tested in the MeWo, 5B1, MCF-7 cell lines for analyzing CD98hc, ST3GAL1 and ST3GAL2 protein levels. For all Western blot analysis using mimics, miR-548ab was used as the non-targeting control (NTC) based on experimental data (see Supplementary Fig. 4).

For all experiments, cells were seeded in six-well plates (50,000 cells/well) and cultured for 24 h in appropriate media. Cells were then washed with Hanks buffered salt solution (HBSS, Gibco) and transfected with miRNA mimics: 50 nM mimic (Dharmacon, Horizon Discovery), 5 μl Lipofectamine 2000 (Life Technologies), 250 μl OptiMEM. The media was changed to standard media 12 h post-transfection. Cells were then lysed at 48 h post-transfection in cold RIPA buffer supplemented with Halt™ protease inhibitors (Thermofisher, catalog #: 89900). For Western blot analysis, 50 µg of protein was added to 4x loading buffer with DTT (1 mM), heated at 95 °C for 10 min and run on a 10% gel (SDS-PAGE) using standard conditions. Proteins were then transferred from the gel to Polyvinylidene fluoride membrane using iBlot2 Transfer Stacks (PVDF, Invitrogen, catalog number: IB24002) and the iBlot2 transfer device (Invitrogen) using the standard protocol (P_0_). Blots were then incubated with Ponceau S Solution (Boston BioProdcuts, catalog #ST-180) for 3 min and the total protein levels were imaged using the protein gel mode (Azure 600, Azure Biosystems Inc.). Blots were blocked with 5% BSA (CD98hc and ST3GAL1) or 5% non-fat dry milk (ST3GAL2) in TBST buffer (TBS buffer plus 0.1% Tween 20) for 30 min at room temperature on rocker (LSE platform rocker, Corning) with 60 rpm. For CD98hc, blots were then incubated with Monoclonal mouse α-human-CD98hc 1° antibody (1:500 in TBST with 10% BSA, catalog #: 66883-1-IG, Proteintech). For ST3GAL1, rabbit α-human-ST3GAL1 1° antibody (1:1000 in TBST with 10% BSA, catalog #: HPA040466, Sigma) was used. For ST3GAL2, rabbit α-human-ST3GAL2 1° antibody (1:1000 in TBST with 10% non-fat dry milk, catalog #: ab96028, Abcam) was used. After an overnight incubation at 4 ℃, blots were washed 3 × 1 min with 0.1% TBST buffer. A secondary antibody was then added (ST3GAL1/2:α-rabbit IgG-HRP, 1: 6,000 in TBST with 10% BSA (ST3GAL1) or non-fat dry milk (ST3GAL2); CD98: α-mouse IgG-HRP, 1: 5,000 in TBST with 10% BSA) and incubated for 1 h at room temperature with shaking (60 rpm). Blots were then washed 3 × 1 min with 0.1% TBST buffer. Blots were developed using Clarity and Clarity Max Western ECL substrate according to the manufacturer’s instructions (Bio-Rad). Membranes were imaged in chemiluminescent mode (Azure 600, Azure Biosystems Inc.). All analysis was done in a minimum of three biological replicates.

### Endogenous miRNA Activity Validation

miRIDIAN microRNA Hairpin Inhibitors (CD98hc: anti-miRs -155-5p, -135b-5p; ST3GAL1: anti-miRs -1246, -28-3p, -449b-5p; ST3GAL2: anti-miRs -30b-5p, -30c-1-3p, - 378j) and miRIDIAN microRNA Hairpin Inhibitor Negative Control (NTC) were purchased from Dharmacon (Horizon Discovery, Cambridge, UK). The selected anti-up-miRs for each protein were tested in different cell lines: CD98hc: MeWo and 5B1 cells; ST3GAL1: SK-OV-3 cells; ST3GAL2: A375 cells. For co-up-miRs (CD98hc/ST3GAL1: anti-miRs - 1246, -28-3p, -449b-5p; CD98hc/ST3GAL2: anti-miRs -30b-5p, -30c-1-3p, -378j) were tested in the MeWo, 5B1 cell line for analyzing CD98hc, ST3GAL1 and ST3GAL2 protein levels. All cell lines were seeded and incubated as described for Western blot analysis. Cells were transfected with anti-miRNAs, 50 nM using Lipofectamine™ 2000 transfection reagent in OptiMEM following the manufacturer’s instructions (Life Technologies) as described above. After 12 h media was changed to standard culture media. 48 h post-transfection cells were lysed and analyzed for CD98hc, ST3GAL1 and ST3GAL2 protein levels as previously described. All analysis was done in a minimum of three biological replicate.

### RT-qPCR

Total RNA was isolated from cells treated as in Western blot experiments using TRIzol reagent (catalog #: 15596018, Invitrogen) according to the manufacturer’s instructions. RNA concentrations were measured using NanoDrop. Isolated total RNA (1 μg) was then reverse transcribed to cDNA using Superscript III Cells Direct cDNA synthesis kit (catalog #: 18080300, Invitrogen). Reverse transcription quantitative PCR (RT-qPCR) was performed using the SYBR Green method and cycle threshold values (Ct) were obtained using an Applied Biosystem (ABI) 7500 Real-Time PCR machine. Values were normalized to the housekeeping gene GAPDH. The primer sequences used for RT-qPCR can be found in Supplementary Table 1. All analysis was done in three biological replicates.

### Labeling of diCBM40

diCBM40 protein was expressed and purified as previously described ^60^. Protein was fluorescently labelled with Cy3- or Cy5-NHS ester (catalog #: GEPA13101 or GEPA15100, Sigma) as follows. First, 2 mg of total protein was diluted in 1X PBS to 1.78 mL. The pH of the solution was adjusted with 200 μl of 1M sodium bicarbonate, then 20 μl of a stock solution (10 mg/mL) of Cy3 or Cy-5 NHS ester was added to the mixture. The mixture was reacted for 1 hour in the dark at room temperature while shaking. Unconjugated dye molecules were then removed by a Zeba™ Dye and Biotin Removal Spin Column (catalog #: A44301, Thermofisher Scientific). The Cy5-diCBM40 was used in flow cytometry experiments amd Cy3-diCBM40 was used in microscopy experiments as described below.

### Fluorescence microscopy

Cells were seeded onto sterile 22 × 22 glass coverslips placed into 35 mm dishes at a density of 2 × 10^5^ cells/ml in the appropriate media for the cell line (see above). After 24h, cells were transfected with miRNA mimics as previously described. At 48 h post-transfection, cells were washed with TBS (3 × 2 mL) and fixed with 4 % paraformaldehyde in PBS for 20 min. Cells were again washed with TBS (3 × 2 mL) prior to incubation with Cy3-diCBM40 (2 mL total volume, 20 μg lectin in TBS, 0.1% BSA) for 1 hour at room temperature in the dark.

Coverslips were then washed (TBS, 3 x 2 mL), and cells were counterstained with Hoechst 33342 (2 mL, 1 μg/mL in TBS, 15 min in a 37 °C incubator). The coverslips were then mounted onto slides with 60 μl of mounting media (90% glycerol in PBS) and imaged with a Zeiss Axio Observer inverted fluorescence microscope (Camera: Axiocam 305 mono, software: ZEN 3.2 pro). Specificity of diCBM40 staining was confirmed by using Neuraminidase (catalog #: P0722L, NEB) prior to Cy3-diCBM40 staining (Supplementary Figs. 11 & 15). All analysis was done in three biological replicates.

### Flow cytometry

*Lectin staining and flow cytometry analysis*: miRNA or anti-miR transfection of cell lines for flow cytometry analysis followed the same method as for Western blot analysis. After 48 h, samples were trypsin digested (100 µl of 0.25 % trypsin per well in 6-well plate format). Up to 1 mL of 1X HBSS was added to remove all the cells from the flask, and cells were pelleted out by centrifugation at 350 x g for 6 min. Cell pellet was resuspended in 1X TBS buffer containing 0.1% BSA and were counted using the cell counter. 100 µl of 5 x 10^5^ cells were used per sample. As a negative control for lectin staining, neuraminidase (catalog #: P0722L, NEB) treated cells were used. Samples were treated with 15 μg/ml of Cy5-diCBM40 in 1X TBS buffer and incubated for 25 min at room temperature in the dark. Cells were then pelleted by centrifugation ( 350 x g, 6 min) and washed 2x with 1X TBS, 0.1%BSA. Samples were then resuspended in 400 µl FACS buffer (PBS, 0.1% BSA, 0.1% EDTA, 5 mM) and analyzed by flow cytometry (see below).

*Cell surface CD98hc staining*: Samples were prepared as above with the following exception: instead of lectin, samples were incubated with FITC labeled anti-CD98 antibody (ab26010, Abcam) at 4 μg/ml (100 μL total volume in TBS, 0.1% BSA). FITC labeled mouse monoclonal IgG1 (ab91356, Abcam) was used as an isotype control for this antibody. Samples were resuspended in 400 µl FACS buffer (PBS, 0.1% BSA, 0.1% EDTA, 5 mM).

*Flow cytometry analysis:* All samples were analyzed using a Katz Fortessa flow cytometer (core facility, Faculty of Medicine and Dentistry, University of Alberta) and 20,000 cells were analyzed per sample. All experiments are done in a minimum of three biological replicates. The data was analyzed using FlowJo (10.5.3) software (BD Biosciences).

### Multi-Site Mutagenesis on pFmiR-ST3GAL1, pFmiR-ST3GAL2, pFmiR-CD98hc

The 3’UTR sequence of CD98hc, ST3GAL1, ST3GAL2 and the corresponding miRNA sequences (CD98hc: miRs -1246, -30b-5p, -155-5p, -28-3p; ST3GAL1: miR-1246; ST3GAL2: miR-30b-5p) were analyzed using RNAhybrid ^53^ which calculates a minimal free energy hybridization of target RNA sequence and miRNA. The most stable predicted miRNA:mRNA interaction sites were selected and mutant pFmiR-sensors designed. Mutatation primers (IDT DNA) were designed using NEB Base Changer version 1 (https://nebasechangerv1.neb.com) and are listed in Supplementary Table 1. Multiple mutation sites were achieved using Site Directed Mutagenesis (SDM) with the Q5^®^ Site-Directed Mutagenesis kit (NEB, catalog #: E0554S). Amplicons were cleaned up using Monarch^®^ PCR & DNA cleanup kit (catalog #: T1030S, NEB). Sequences for the mutant pFmiR-ST3GAL1, pFmiR-ST3GAL2, pFmiR-CD98hc sensors were verified by Sanger sequencing and used in the miRFluR assay as described previously. A minimum of 5-wells were transfected per sensor and the analysis was done in three biological experiments. Data was normalized to miR-548ab as the NTC.

## Acknowledgements

Funding for L.K.M. comes from the Canada Excellence Research Chairs Program (CERC in Glycomics). Experiments were performed at the University of Alberta Faculty of Medicine & Dentistry Flow Cytometry Facility, RRID:SCR_019195, which receives financial support from the Faculty of Medicine & Dentistry and Canada Foundation for Innovation (CFI) awards to contributing investigators. Figures 1, 2, & 3 were partially made using BioRender.

## References

1 Uhlen, M. et al. Proteomics. Tissue-based map of the human proteome. Science 347, 1260419 (2015). 10.1126/science.1260419

2 Lundberg, E. et al. Defining the transcriptome and proteome in three functionally different human cell lines. Mol Syst Biol 6, 450 (2010). 10.1038/msb.2010.106

3 Bartel, D. P. Metazoan MicroRNAs. Cell 173, 20–51 (2018). 10.1016/j.cell.2018.03.006

4 Schmiedel, J. M. et al. Gene expression. MicroRNA control of protein expression noise. Science 348, 128–132 (2015). 10.1126/science.aaa1738

5 Thu, C. T., Chung, J. Y., Dhawan, D., Vaiana, C. A. & Mahal, L. K. High-Throughput miRFluR Platform Identifies miRNA Regulating B3GLCT That Predict Peters’ Plus Syndrome Phenotype, Supporting the miRNA Proxy Hypothesis. ACS Chem Biol 16, 1900–1907 (2021). 10.1021/acschembio.1c00247

6 Jame-Chenarboo, F., Ng, H. H., Macdonald, D. & Mahal, L. K. High-Throughput Analysis Reveals miRNA Upregulating alpha-2,6-Sialic Acid through Direct miRNA-mRNA Interactions. ACS Cent Sci 8, 1527–1536 (2022). 10.1021/acscentsci.2c00748

7 Vasudevan, S. & Steitz, J. A. AU-rich-element-mediated upregulation of translation by FXR1 and Argonaute 2. Cell 128, 1105–1118 (2007). 10.1016/j.cell.2007.01.038

8 Vasudevan, S., Tong, Y. & Steitz, J. A. Switching from repression to activation: microRNAs can up-regulate translation. Science 318, 1931–1934 (2007). 10.1126/science.1149460

9 Zhang, X. et al. MicroRNA directly enhances mitochondrial translation during muscle differentiation. Cell 158, 607–619 (2014). 10.1016/j.cell.2014.05.047

10 Eulalio, A., Tritschler, F. & Izaurralde, E. The GW182 protein family in animal cells: new insights into domains required for miRNA-mediated gene silencing. RNA 15, 1433–1442 (2009). 10.1261/rna.1703809

11 Agrawal, P., Chen, S., Pablos, A.D., Jame-Chenarboo, F.,Vega, E.M.S.D.,Darvishian, F.,Osman, I.,Lujambio, A.,Mahal, L.K., Hernando, E. Integrated in vivo functional screens and multi-omics analyses identify a-2,3-sialylation as essential for melanoma maintenance. (2024). 10.1101/2024.03.08.584072

12 Fenczik, C. A. et al. Distinct domains of CD98hc regulate integrins and amino acid transport. J Biol Chem 276, 8746–8752 (2001). 10.1074/jbc.M011239200

13 Ip, H. & Sethi, T. CD98 signals controlling tumorigenesis. Int J Biochem Cell Biol 81, 148–150 (2016). 10.1016/j.biocel.2016.11.005

14 Cantor, J. et al. CD98hc facilitates B cell proliferation and adaptive humoral immunity. Nat Immunol 10, 412–419 (2009). 10.1038/ni.1712

15 Cantor, J., Slepak, M., Ege, N., Chang, J. T. & Ginsberg, M. H. Loss of T cell CD98 H chain specifically ablates T cell clonal expansion and protects from autoimmunity. J Immunol 187, 851–860 (2011). 10.4049/jimmunol.1100002

16 Cantor, J. M. & Ginsberg, M. H. CD98 at the crossroads of adaptive immunity and cancer. J Cell Sci 125, 1373–1382 (2012). 10.1242/jcs.096040

17 Yagita, H., Masuko, T. & Hashimoto, Y. Inhibition of tumor cell growth in vitro by murine monoclonal antibodies that recognize a proliferation-associated cell surface antigen system in rats and humans. Cancer Res 46, 1478–1484 (1986).

18 Ichinoe, M. et al. Prognostic values of L-type amino acid transporter 1 and CD98hc expression in breast cancer. J Clin Pathol 74, 589–595 (2021). 10.1136/jclinpath-2020-206457

19 Kaira, K. et al. CD98 expression is associated with poor prognosis in resected non-small-cell lung cancer with lymph node metastases. Ann Surg Oncol 16, 3473–3481 (2009). 10.1245/s10434-009-0685-0

20 Theodosakis, N. et al. Integrative discovery of CD98 as a melanoma biomarker. Pigment Cell Melanoma Res 29, 385–387 (2016). 10.1111/pcmr.12464

21 Bajaj, J. et al. CD98-Mediated Adhesive Signaling Enables the Establishment and Propagation of Acute Myelogenous Leukemia. Cancer Cell 30, 792–805 (2016). 10.1016/j.ccell.2016.10.003

22 Chandrasekaran, E. V. et al. Mammalian sialyltransferase ST3Gal-II: its exchange sialylation catalytic properties allow labeling of sialyl residues in mucin-type sialylated glycoproteins and specific gangliosides. Biochemistry 50, 9475–9487 (2011). 10.1021/bi200301w

23 Priatel, J. J. et al. The ST3Gal-I sialyltransferase controls CD8+ T lymphocyte homeostasis by modulating O-glycan biosynthesis. Immunity 12, 273–283 (2000). 10.1016/s1074-7613(00)80180-6

24 Dalziel, M. et al. The relative activities of the C2GnT1 and ST3Gal-I glycosyltransferases determine O-glycan structure and expression of a tumor-associated epitope on MUC1. J Biol Chem 276, 11007–11015 (2001). 10.1074/jbc.M006523200

25 Walker, M. R. et al. O-linked alpha2,3 sialylation defines stem cell populations in breast cancer. Sci Adv 8, eabj9513 (2022). 10.1126/sciadv.abj9513

26 Wu, X. et al. Sialyltransferase ST3GAL1 promotes cell migration, invasion, and TGF-beta1-induced EMT and confers paclitaxel resistance in ovarian cancer. Cell Death Dis 9, 1102 (2018). 10.1038/s41419-018-1101-0

27 Pietrobono, S. et al. ST3GAL1 is a target of the SOX2-GLI1 transcriptional complex and promotes melanoma metastasis through AXL. Nat Commun 11, 5865 (2020). 10.1038/s41467-020-19575-2

28 Hong, Y. et al. ST3GAL1 and betaII-spectrin pathways control CAR T cell migration to target tumors. Nat Immunol 24, 1007–1019 (2023). 10.1038/s41590-023-01498-x

29 Deschuyter, M. et al. ST3GAL2 knock-down decreases tumoral character of colorectal cancer cells in vitro and in vivo. Am J Cancer Res 12, 280–302 (2022).

30 Lin, C. W. et al. Homogeneous antibody and CAR-T cells with improved effector functions targeting SSEA-4 glycan on pancreatic cancer. Proc Natl Acad Sci U S A 118 (2021). 10.1073/pnas.2114774118

31 Gaziel-Sovran, A. et al. miR-30b/30d regulation of GalNAc transferases enhances invasion and immunosuppression during metastasis. Cancer Cell 20, 104–118 (2011). 10.1016/j.ccr.2011.05.027

32 Gómez-Martínez, H., López, A., Gil, M.P., Hernando, E., García-García, F. Novel diagnosis and progression microRNA signatures in melanoma. (2023). 10.1101/2023.10.20.563284

33 Xia, P. & Dubrovska, A. CD98 heavy chain as a prognostic biomarker and target for cancer treatment. Front Oncol 13, 1251100 (2023). 10.3389/fonc.2023.1251100

34 Ekiz, H. A. et al. MicroRNA-155 coordinates the immunological landscape within murine melanoma and correlates with immunity in human cancers. JCI Insight 4 (2019). 10.1172/jci.insight.126543

35 Peng, J., Liu, H. & Liu, C. MiR-155 Promotes Uveal Melanoma Cell Proliferation and Invasion by Regulating NDFIP1 Expression. Technol Cancer Res Treat 16, 1160–1167 (2017). 10.1177/1533034617737923

36 Hua, K. et al. miR-135b, upregulated in breast cancer, promotes cell growth and disrupts the cell cycle by regulating LATS2. Int J Oncol 48, 1997–2006 (2016). 10.3892/ijo.2016.3405

37 Hossain, A., Kuo, M. T. & Saunders, G. F. Mir-17-5p regulates breast cancer cell proliferation by inhibiting translation of AIB1 mRNA. Mol Cell Biol 26, 8191–8201 (2006). 10.1128/MCB.00242-06

38 Cruz-Munoz, W., Man, S., Xu, P. & Kerbel, R. S. Development of a preclinical model of spontaneous human melanoma central nervous system metastasis. Cancer Res 68, 4500–4505 (2008). 10.1158/0008-5472.CAN-08-0041

39 Barshir, R. et al. GeneCaRNA: A Comprehensive Gene-centric Database of Human Non-coding RNAs in the GeneCards Suite. J Mol Biol 433, 166913 (2021). 10.1016/j.jmb.2021.166913

40 Kurcon, T. et al. miRNA proxy approach reveals hidden functions of glycosylation. Proc Natl Acad Sci U S A 112, 7327–7332 (2015). 10.1073/pnas.1502076112

41 Buccitelli, C. & Selbach, M. mRNAs, proteins and the emerging principles of gene expression control. Nat Rev Genet 21, 630–644 (2020). 10.1038/s41576-020-0258-4

42 Vogel, C. & Marcotte, E. M. Insights into the regulation of protein abundance from proteomic and transcriptomic analyses. Nat Rev Genet 13, 227–232 (2012). 10.1038/nrg3185

43 Stinson, S. et al. TRPS1 targeting by miR-221/222 promotes the epithelial-to-mesenchymal transition in breast cancer. Sci Signal 4, ra41 (2011). 10.1126/scisignal.2001538

44 Kong, W. et al. Upregulation of miRNA-155 promotes tumour angiogenesis by targeting VHL and is associated with poor prognosis and triple-negative breast cancer. Oncogene 33, 679–689 (2014). 10.1038/onc.2012.636

45 Lee, E. J. et al. Expression profiling identifies microRNA signature in pancreatic cancer. Int J Cancer 120, 1046–1054 (2007). 10.1002/ijc.22394

46 Bayraktar, R. & Van Roosbroeck, K. miR-155 in cancer drug resistance and as target for miRNA-based therapeutics. Cancer Metastasis Rev 37, 33–44 (2018). 10.1007/s10555-017-9724-7

47 Moutabian, H. et al. MicroRNA-155 and cancer metastasis: Regulation of invasion, migration, and epithelial-to-mesenchymal transition. Pathol Res Pract 250, 154789 (2023). 10.1016/j.prp.2023.154789

48 Eis, P. S. et al. Accumulation of miR-155 and BIC RNA in human B cell lymphomas. Proc Natl Acad Sci U S A 102, 3627–3632 (2005). 10.1073/pnas.0500613102

49 Hoang, D. H. et al. MicroRNA networks in FLT3-ITD acute myeloid leukemia. Proc Natl Acad Sci U S A 119, e2112482119 (2022). 10.1073/pnas.2112482119

50 Liu, F., Mao, Q., Zhu, S. & Qiu, J. MicroRNA-155-5p promotes cell proliferation and invasion in lung squamous cell carcinoma through negative regulation of fibroblast growth factor 9 expression. J Thorac Dis 13, 3669–3679 (2021). 10.21037/jtd-21-882

51 Schjenken, J. E. et al. MicroRNA miR-155 is required for expansion of regulatory T cells to mediate robust pregnancy tolerance in mice. Mucosal Immunol 13, 609–625 (2020). 10.1038/s41385-020-0255-0

52 Costinean, S. et al. Pre-B cell proliferation and lymphoblastic leukemia/high-grade lymphoma in E(mu)-miR155 transgenic mice. Proc Natl Acad Sci U S A 103, 7024–7029 (2006). 10.1073/pnas.0602266103

53 Rehmsmeier, M., Steffen, P., Hochsmann, M. & Giegerich, R. Fast and effective prediction of microRNA/target duplexes. RNA 10, 1507–1517 (2004). 10.1261/rna.5248604

54 Kudo, T. et al. Up-regulation of a set of glycosyltransferase genes in human colorectal cancer. Lab Invest 78, 797–811 (1998).

55 Madunic, K. et al. Specific (sialyl-)Lewis core 2 O-glycans differentiate colorectal cancer from healthy colon epithelium. Theranostics 12, 4498–4512 (2022). 10.7150/thno.72818

56 Li, D., Zhong, J., Zhang, G., Lin, L. & Liu, Z. Oncogenic Role and Prognostic Value of MicroRNA-937-3p in Patients with Breast Cancer. Onco Targets Ther 12, 11045–11056 (2019). 10.2147/OTT.S229510

57 Lv, Y., Yang, H., Ma, X. & Wu, G. Strand-specific miR-28-3p and miR-28-5p have differential effects on nasopharyngeal cancer cells proliferation, apoptosis, migration and invasion. Cancer Cell Int 19, 187 (2019). 10.1186/s12935-019-0915-x

58 Song, J. et al. Let-7i-5p inhibits the proliferation and metastasis of colon cancer cells by targeting kallikrein-related peptidase 6. Oncol Rep 40, 1459–1466 (2018). 10.3892/or.2018.6577

59 Wang, X. et al. Circulating exosomal miR-363-5p inhibits lymph node metastasis by downregulating PDGFB and serves as a potential noninvasive biomarker for breast cancer. Mol Oncol 15, 2466–2479 (2021). 10.1002/1878-0261.13029

60 Ribeiro, J. P. et al. Characterization of a high-affinity sialic acid-specific CBM40 from Clostridium perfringens and engineering of a divalent form. Biochem J 473, 2109–2118 (2016). 10.1042/BCJ20160340

61 Yu, Y., Yu, F. & Sun, P. MicroRNA-1246 Promotes Melanoma Progression Through Targeting FOXA2. Onco Targets Ther 13, 1245–1253 (2020). 10.2147/OTT.S234276

62 Komina, A., Palkina, N., Aksenenko, M., Tsyrenzhapova, S. & Ruksha, T. Antiproliferative and Pro-Apoptotic Effects of MiR-4286 Inhibition in Melanoma Cells. PLoS One 11, e0168229 (2016). 10.1371/journal.pone.0168229

63 Haynes, B. F. et al. Characterization of a monoclonal antibody (4F2) that binds to human monocytes and to a subset of activated lymphocytes. J Immunol 126, 1409–1414 (1981).

64 Mayr, C. Evolution and Biological Roles of Alternative 3’UTRs. Trends Cell Biol 26, 227–237 (2016). 10.1016/j.tcb.2015.10.012

65 Mayr, C. Regulation by 3’-Untranslated Regions. Annu Rev Genet 51, 171–194 (2017). 10.1146/annurev-genet-120116-024704

66 Bartel, D. P. MicroRNAs: target recognition and regulatory functions. Cell 136, 215–233 (2009). 10.1016/j.cell.2009.01.002

67 Helwak, A., Kudla, G., Dudnakova, T. & Tollervey, D. Mapping the human miRNA interactome by CLASH reveals frequent noncanonical binding. Cell 153, 654–665 (2013). 10.1016/j.cell.2013.03.043

68 Bartel, D. P. & Chen, C. Z. Micromanagers of gene expression: the potentially widespread influence of metazoan microRNAs. Nat Rev Genet 5, 396–400 (2004). 10.1038/nrg1328

69 Gurtan, A. M. & Sharp, P. A. The role of miRNAs in regulating gene expression networks. J Mol Biol 425, 3582–3600 (2013). 10.1016/j.jmb.2013.03.007

70 Console, L. et al. N-glycosylation is crucial for trafficking and stability of SLC3A2 (CD98). Sci Rep 12, 14570 (2022). 10.1038/s41598-022-18779-4

