## Supplementary Information for "Screening the human miRNA interactome reveals coordinated upregulation in melanoma, adding bidirectional regulation to miRNA networks"

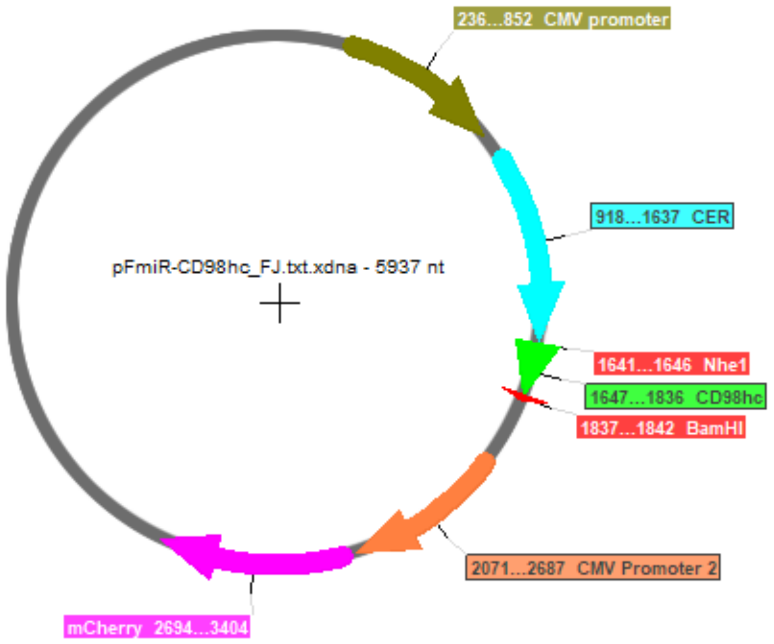


5’CTTCAGCCTGACATGGACCCACTACCCTTCTCCTTTCCTTCCCAGGCCCTTTGGCTTCTGATTTTTCTCTTTTTTAAAAACAAACAAACA
AACTGTTGCAGATTATGAGTGAACCCCCAAATAGGGTGTTTTCTGCCTTCAAATAAAAGTCACCCCTGCATGGTGAAGTCTTCCCTCTGCTT
CTCTCATA3’

**Supplementary Fig. 1. pFmiR-CD98hc map and sequence**. **a**. pFmiR-CD98hc plasmid map. **b**. CD98hc 3'UTR sequence. Sequence contains 190 base pairs.

a

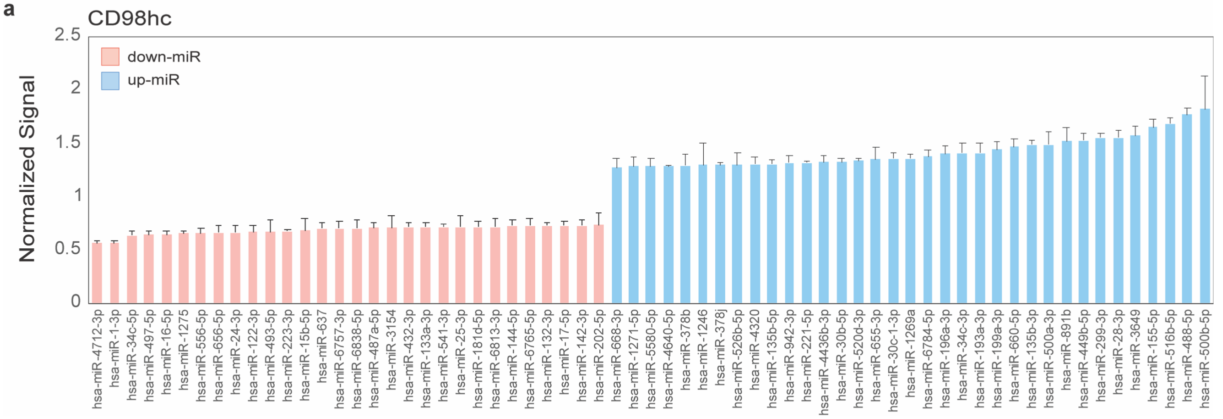


**Supplementary Fig. 2. miRFluR assay hits for CD98hc. a.** Bar graph of miRNA hits from 95% confidence interval for CD98hc. Data are normalized over median. Error bars represent standard deviation of technical replicates (n=3).

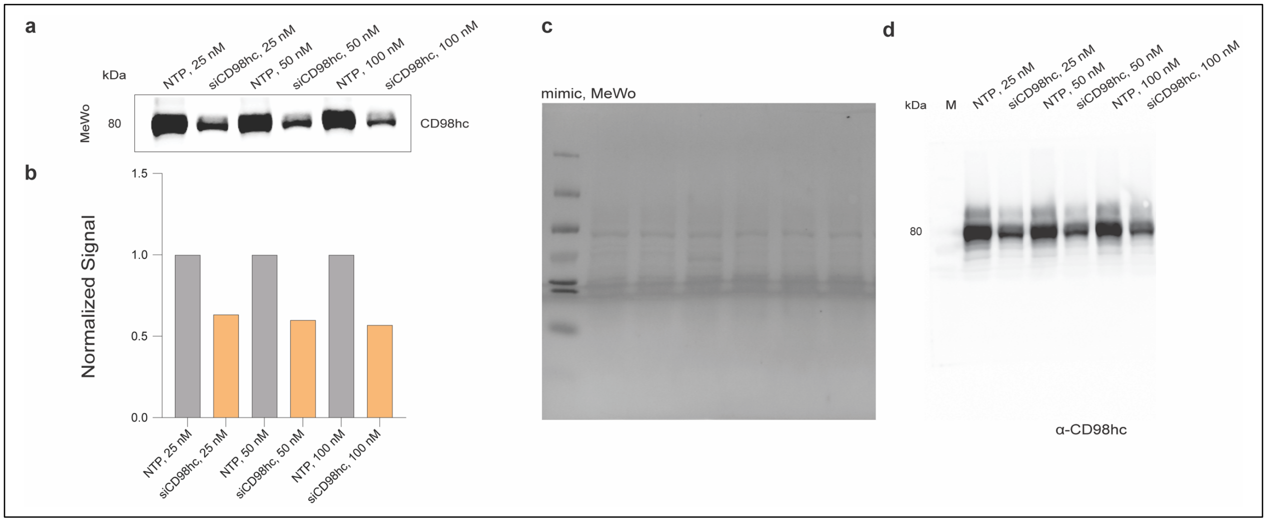


**Supplementary Fig. 3. Validation of antibody for CD98hc. a**. Western blot analysis of MeWo cells treated with non-targeting pooled (NTP) or pooled siRNA targeting CD98hc (siCD98hc). Cells were transfected into with varying amounts of siRNA (25, 50, 100 nM) for 48 h prior to Western blot analysis. **b**. Bar graph shows quantification of results shown in a. Signals were normalized to total protein on Ponceau. **c**. Ponceau of Western blot shown in a. **d**. Complete blot of Western shown in a.

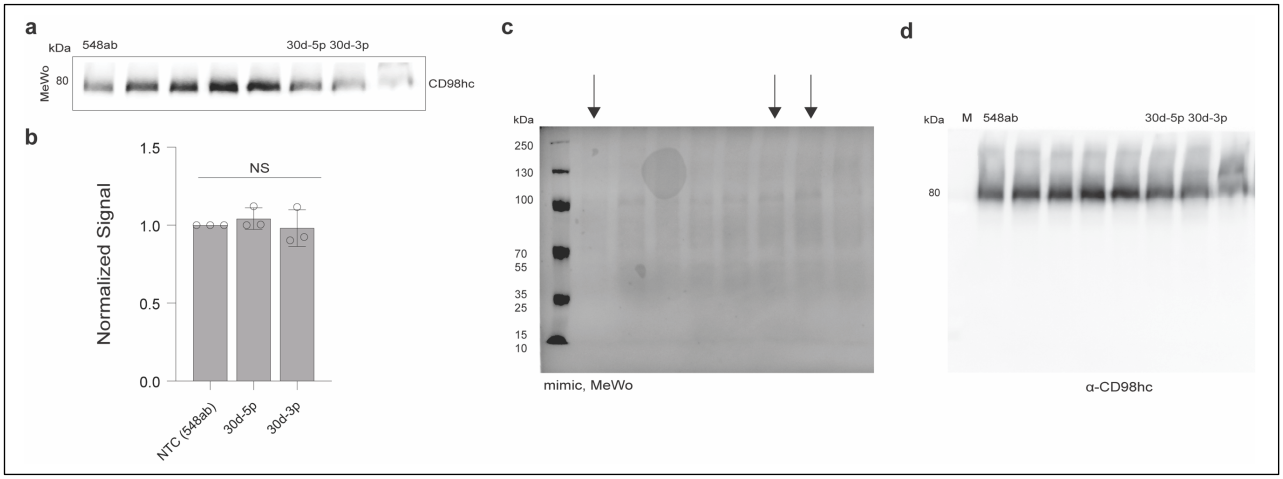


**Supplementary Fig. 4. Validation of miR-548ab as new NTC. a**. miRNA mimics of median controls (miRs: -548ab (NTC), -30d-5p, -30d-3p -indicated on blot) were transfected into MeWo cells (50 nM miR, 48 h) prior to Western blot analysis. Representative data is shown. **b**. Bar graph shows quantification of triplicate Western blot results as obtained in a. Signals were normalized to total protein on Ponceau. **c**. Ponceau of Western blot shown in a. **d**. Complete blot of Western shown in a, median control miRNA are indicated.

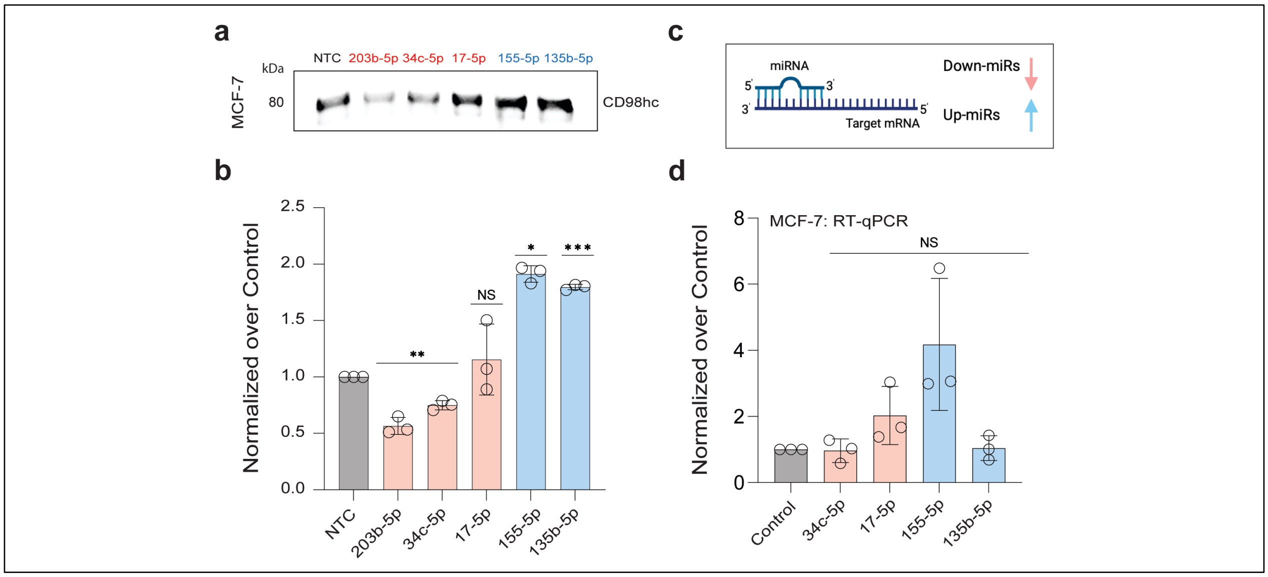


**Supplementary Fig. 5. Validation of miRFluR data for CD98hc in MCF-7 cell line.** Impact of miRNA mimics on CD98hc expression. MCF-7 cells were treated with miRNA mimics (down-miRs: miR-203b-5p, -34c-5p, -17-5p, up-miRs: miR-155-5p, -135b-5p) or non-targeting control (NTC: miR-548ab) at 50 nM for 48h and analyzed as indicated. **a.** Representative Western blot analysis of CD98hc in MCF-7. **b.** Bar graph of Western blot data shown in a. **c**. miRNA: mRNA interaction. **d**. RT-qPCR quantitative analysis of *slc3a2* in MCF-7. Samples were normalized to GAPDH and NTC. The Wilcoxon *t*-test was used to compare miRs to NTC (NS: not significant, * $p < 0.05$, ** $< 0.01$, *** $<0.001$). Western blot and RT-qPCR experiments are performed in independent biological replicates (n=3).

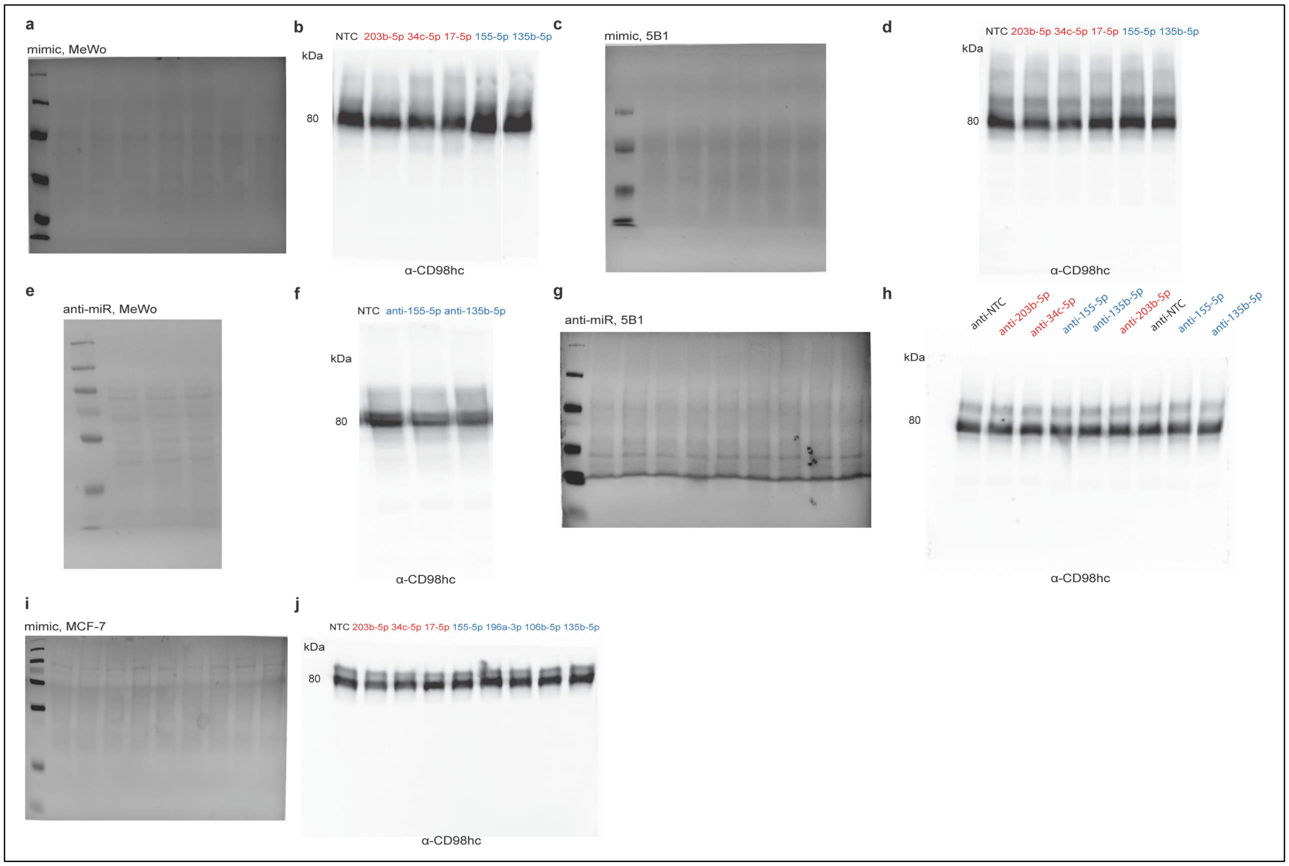


**Supplementary Fig. 6. Ponceau and Whole Western blots for data shown in Fig. 2 and Supplementary Fig. 5. a, c, i.** Ponceau staining of blot used in Fig. 2 (a: MeWo; c: 5B1), and Supplementary Fig. 5 (i: MCF-7). **b, d, j.** Whole Western blot for data shown in Figs. 2 (b: MeWo; d: 5B1), and Supplementary Fig. 5 (j: MCF-7). **e, g.** Ponceau staining of blot used in Fig. 2 (e: MeWo), (g: 5B1). **f, h**. Whole Western blot for data shown in Fig. 2 (f: MeWo; h: 5B1).

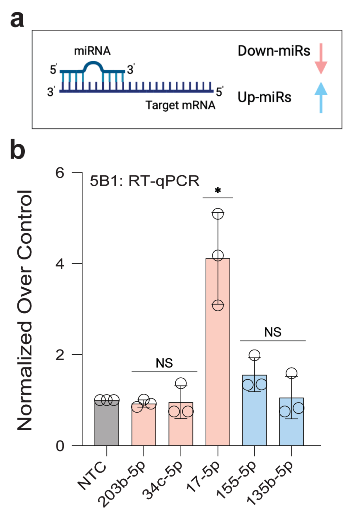


**Supplementary Fig. 7. RT-qPCR in 5B1.** Impact of miRNA mimics on CD98hc expression. 5B1 cells were treated with miRNA mimics (down-miRs: miR-203b-5p, -34c-5p, -17-5p, upmiRs: miR-155-5p, -135b-5p) or NTC at 50 nM for 48h and analyzed as indicated. **a.** miRNA: mRNA interaction. **b**. RT-qPCR analysis for samples as in Fig. 2c-d. Data was normalized to GAPDH and to NTC. The Wilcoxon *t*-test was used to compare miRs to NTC (ns not significant, * $p < 0.05$, ** < *0.01,* *** <*0.001*). RT-qPCR experiment is performed in independent biological replicates (n=3).

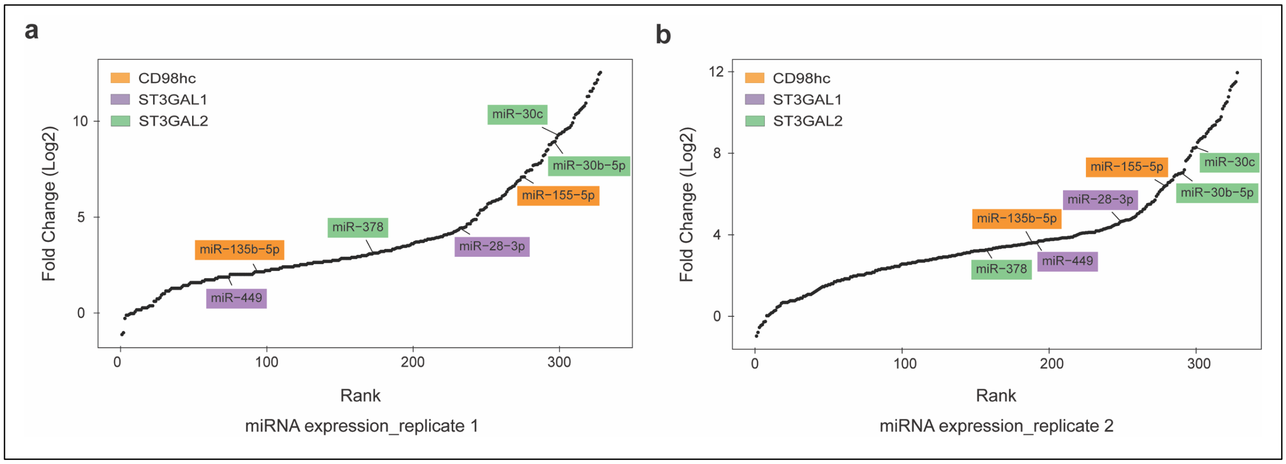


**Supplementary Fig. 8. miRNA expression profiling in MeWo cell line. a, b.** Graphs represent two replicates of miRNA expression profiles in MeWo cell line from GEO dataset (GEO accession number: GSE10833)[34]. miRNA used in this study are highlighted.

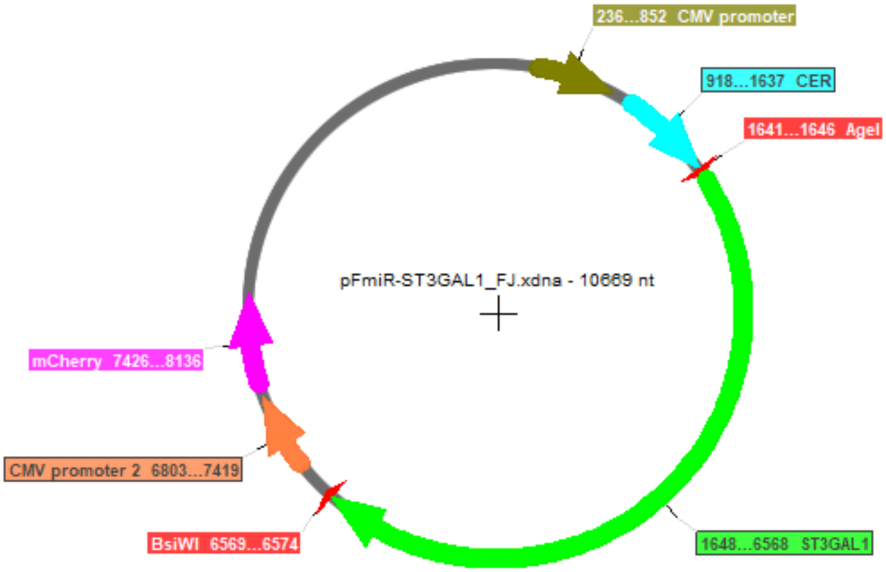


5′CGCAGTGAAGGGCTGAGGATGGACGCACTGTCACACCTCTGCATTTCCAGCCCCAGCATCTTGCTGGAGCCGTTCCATCCCGGAGCTTGGAGGGGCAGCCTCAGGTGTGTGCCTGG
GCACCGCTCACAGCCTCTTGCACCCAGCCGTTGGCAGCATCTACTCAGCAAGGTCACTAAGCTCTGCCAGCGTGGCAGAGCATGTCTTGGAACCTGTCTTGAGTGGGGACAACGTCCC
CCCACTGCTGCCCTAGAGCTGGGGAGACGCTGGGAAAGGTTCAACCTCCACACACTAAAATCATTTTGGCTCCTGGGGCAAGCTTGGGGAATGAATGTGGAAGATGCCTATATTCTGA
GAGACAGGACAGTTTCCCAGGAAGATGGGCAGAGACTTGAGTGGCGATTACCTCCAGCACAGAGACGTGCCAGGCGGTGTTGGCGCTCGGGGCGAGATGCTGCCCTTCTTTGCACGAA
GCCTGGCCTCTTGCTTGGCGTGATAACCCTGTCATCTTCCCAAAGCTCATTTATGAGCCACCAGAGGCTCCTACCCCAAAGATTTTCACAGAAACTTGAGGCCAGGTGCCGTGGCTCA
CACCTGTAATCTGAACACTTTGGGAGGCCGAGGCGGGAGGATCACTTGAGCCCAGGAGTTCAAGACCAGCCTGGGCAACATAGTGAGACTCCTGTCTCTACAAAAATAAAAGATTTAA
AAAAATTAGCCAGGCACGGTGGCACACACTTGTAGCCCCAGCTACTAGGGAGGCTGAGGAGGGAGGATCTCTTGTGCCTAGGAGTTCGAGGCTGCAGTGGGCTGTGATCACACCACTG
CACTCCAGCCTGGGCAACAGAGTGAGACCCTGTCTCTGAAAAAAAAAAAAAAAAGAAAAGAAAAAAAAACCCCAAACTTGGCACCGTGTTTCTGACTCCTGCAGATGCCTCTTCCCAAGC
AGTTGTTCCTTTGAGTAGTCAGTCTCATGGTTCTAATCGTGGCGAGAGCGGATCTGACCACTTCCATGTGCCTTTGTATTTGGGAAAGAAAAGCATGCATTGGCTGAAATGAAACAAA
AGCCCTTACTTGGGAAAAGGGACCCCCACCAGCGGCCCTGCTTGCTCCGTAGCTGGGAACATCTTTTCTGGTTTCCTGAGCCAGCACCTGAATTGCAACGAAGCTCTGAGGATACTCA
AACCACCCAGCAGTGCGTCAGAATGGAACGTACATTCTCATTCTCAGACTGTTCAATATGGTGCCCGTAACCTCCAAAGCCACACAGGAGGCTGCCAGCAGCATTTTCCCCAAGGGCC
TGAGCTCAGGGTCAAGTCCCTGCTGGCCATCTTGGGGCTGTCACTGGGGGTCTGGGTCAAGGACATCCCAGGAATGGTAACTGCCACATACCTCATCTTCTTACTCCCCAGCTACTCT
CTTTTCCTCATTAGCCTTCCTGATTATTTATTGATTTCTTTTTTTTATTCAACTGACATTGTTTGCACCCTGGCTCTGTTGGTGGTCCTGGTGTTTTAATGAAGCTTGCCTCACCCAG
TAGCTGGGAAATTAGGTGGGGGTGCTTAAGGGAGGTTTGTAGGAGTCCTCGGACGTGGGATATTGGGGGTATCTAGGGCTTGGATGGTTTGTGGAATGTCCTCAGCATCCTCCCGTCC
CAACCCAACCTTCCTTCTTCCTCATTGTGGACTTGGATTTATTCTCAGGCTAGGAGTCTCCAAAACTCCGTGAAACCTTATTGCTGCTGGGTGTGGAGTGGAGAGTGACAGGAATGTG
GGGGACGGTGATTGTTGGCTCCAGTTACTAATGTTTAAGCAGAAAAACCACTTTCTTGTTGAAAATATCCCTGAAAAGGTCGCTGCTGCTTTGAGGAGGCGTGACTTTTTCCTGGTGG
AGTGTGACGGAGGCATGGGGCCTGTGCTCCCAGGCTCGGGGGAGCCCCCTCTGCACACGGGGAGGGAGGAACTTGGGGACATCAGTCATGGGTCACAGGTGGCTTCAGGCCATTCACA
AGAGTCCCCGGAAGTTGCGTGCCAAGTTTGGTTTTTTCCCTGAATCTCTGCATATGAGCATTTTTCTGGAGAAAGGACCATAGCTTTTATTAGATTCTCAAAGGTTCAGAATGGGGAA
TGTGTTAAACAAACCGAAAACCTGTTTCAGCTTTAAAATCCTCCAAAACTTTAAAGACTTTTCTTTTTTCTGAGAAAGGCTGGTAAATCACTGAGTCATCATTCAGAACATACAGCCT
CGAAGGGCAGAGCTCACACCCATATATGACGTTAAGAGCTCAACAAATGTTTGTTGAGTGACTGAGTGACCCCAGGGGCAGACTCCAGATCATCAGACACTGATACAGAGAGCGATGC
TATCTCCTCAACTCTCCCTCTTCTCCTAAAAGCTGAGTCTATATAGGGATGACAGGCTCAGGCTGAGCAGTCGACTTCTCTGGGTTTTGCTGTGTAAGAATAGGTCCCTCAACATGAA
GATGTGTCTGCGTCTGAGCCAGATGCTGCCTGGGTCGGTTGCCTGGCCTCCGTCTGGTCTCCCTCCCTCCTACTAGGGTGGTCAAGCTGTGAGATCTCTTCCAGCCTCCAAATCCAGC
TTTCCCATTTTACTGATGAGAAAATCAAGGCCACATTGCAGCTGGGTGGGGAGTGGAGCGTGACAGGCACGGGAGAGGACAGCGATTGTTGACTCTAGTTCCTGATGTTTAATCAGAA
AAACCACTTTTCCTGTAGAGCACATTTCCTAAAAGGCTGCTGCTGTGTAGGGAGGTGTGGCCTTTTCCTGGTGCAGTGTGAAGGAGGGAAGTGACGGGCAGTCGTGAACTGTCTGGGA
CAGATAGCACAAGTCAGAGTTTGAGCAAAACTAGGGTGTTACCGGCGCTATGTTGAAATTGTCTGGAGGCGGGCTTTCCCAGCGAGTCTCCATTCAGCATCTGGAAAAAGTGTAACCA
GAATGGAGGTGTCTGACTCTCCTGTGCTCTCTCAGCGCCGAGCTCTAGCTGCAGGCTGCCTGGGTGGCCTTCTCACACCCTCCCGTGCAGGAGACTGCATTTCTCTTTCCCACAGCCC
AGCAGATAGCTCATTGGCTTAGCAATGGGCACTTAACAGAATCCAGTTGTTGCAGCCAGAATTAAATGCACCAGTCTGGCTTAAACCAACTGGAACTCATCCCTGGGGTGAAGGTGGC
TGGGAATTAGACAAGAGTGTGTGTTAGCCAAAAGGCACATGCATGCTGTCAGCAGGAGAATGAGGAGCACTGGTTCTGGGCAGGCAGTGGCAGGGGCTCCCTCAGGCCCCTGATTCCC
AGCCCACAGCTCTTTCTACTCTGCAGACAGAGTTCCTTGGAGACTGGCCCTGCATTCAGGCAGGGAGCAGAGGGGTCCCTGCCTGCAGCACCCAGAGCTGAACCTGAAGCTACAAAGA
TGGGCTGATGAGGTGGCCCATGAGGTGTCAGCAACATCCCAACCCCCTGGGTCTCCCCAGTGAGTCAGGGCACTGATTTGCTATAGCTGGAGCTACATGCACCTCTCCTCCCCTCAGC
CACCTTTTAGGGGAGAAAGGAGGGCAGATTTGTAACCTCTGGTTCTCTGCAGGGATCCTGGGCTGGAAGAAAGTTCATAAATGGATGAATGCCTTTTAAAACTCCAGCTTTTCCAAGC
ATCATTCACCAAAGCATTTGGTTAGGCTTAGAGAGAAAAGCTGGATATGAGAAAATAGAATGTAAAATAGGAGGGAAAAAAAAAAAGAAGGTGGAGAGGGCAGTTAAGATCTGGGTGA
TTCTTTAGGGAAACGGCTGGTTGAGAGGCTTTGATGGAATCCCATTCTCTGCTGTGGCCAGAAGGTGGTGCTAGAGGCCAGTCCCACTTGGATGTGACCCTGGCCTCTGCCAGGATTC
AAACCACAAAGAACATTTGGAACTGGGATGTTTGCTTTGAAGCAGCCCGAAAACAAGTGAAAGGAAACCAAGATAACTCTCTCTCCCTCCGCTGGCAAGGATCAAAAATACCTTGTGC
AGGGAAGCCCGAGCCTCGAGGGGCCACTGCCCAGCTAAAGATGACTGGACACCAGCCATGCACCATCAGGAACCAGCACCTGTCCTCACCCCTGAGAAGAGCCCCGGGGGTATCATTA
TATCCATCGCAGACATCTGGTTTGCAGCAAATGTGGCTACACGCCATCAGGATTATCCTGCCCTGTGGACTGGGCTAGCTAGTTTTGAGATAGAGTCAACCTCCCCATTTTTGTTTGT
CATGAAAACACCAGCTCCTGCACCCCTGGGGTGCTACCTGACACCTGTGCAAAGAAAAGGAGGAAAACTTAGAAGAAAAGGCAGAGACAGGCAAGACAGCATCTCACAGCTTGTCAGG
ATCAGGGCATTACCGACTATTTTTTCTTACCAAAAGAGCACTGAATTTCCTAGGGGAAGGTGAGCCACCTTTACACCTCCTGTCCTCCCTCCCAGCCCGTCAAGCGATCCCATCACCC
AGCCTCATCACTCAGCTTCTTCCTAGTACAAAACTGCCATCGGTTGTACGTCCTGAGCTGTGGGACGTCAGCTTTAACCCTAGCCTGTCTCAGGGCCTCGCCCTCCAGCCCCCCGCCC
CTTAGGGTGATCGTGGGTTTCTTCCCTTCCTCCTGCACAATCGCTTCCCTTCACTGAATTCCCTTCAAAGCCAAAGTTTCTTGAGCACTTTTTCTTGTGTGCAGTCTTGTGTGTTTGG
GTGTTGGTGTTTTTAAATTTGCTTTCTATTCCCCCCTGCCTCTCTCCATGTCGTTACGCAGCTGAGCACTGGTTGGGATGCATTCGAATGACTTCAGCAAGATCCTGCTTTCCTCAGC
CCCCGCTCTGTTTTATTTATTGTTTAATATTTCCAATGATCCAAATCAAAGTGAATAAACAAAGAGCTATTTTATTGTT3′

**Supplementary Fig. 9. pFmiR-ST3GAL1**. **a**. pFmiR-ST3GAL1 plasmid map. **b**. Sequence of ST3GAL1 3'UTR (4916 bp).

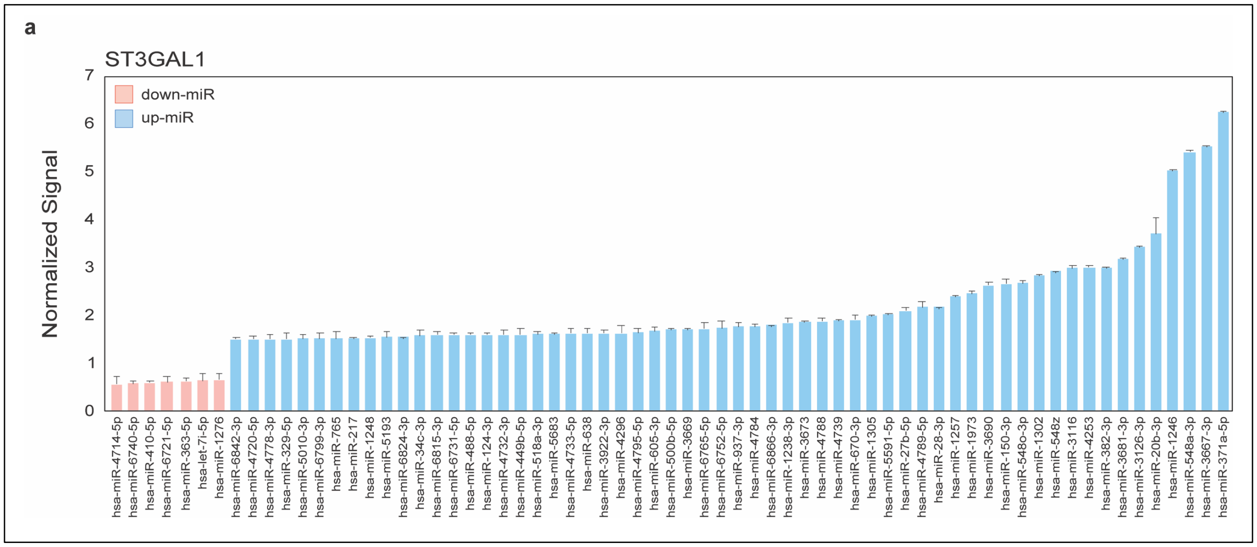


**Supplementary Fig. 10. Hits for ST3GAL1 in miRFluR assay**. **a**. Bar graph of miRNA hits (95% confidence interval) for ST3GAL1. Data are normalized over median. Error bars represent standard deviation of technical triplicates.

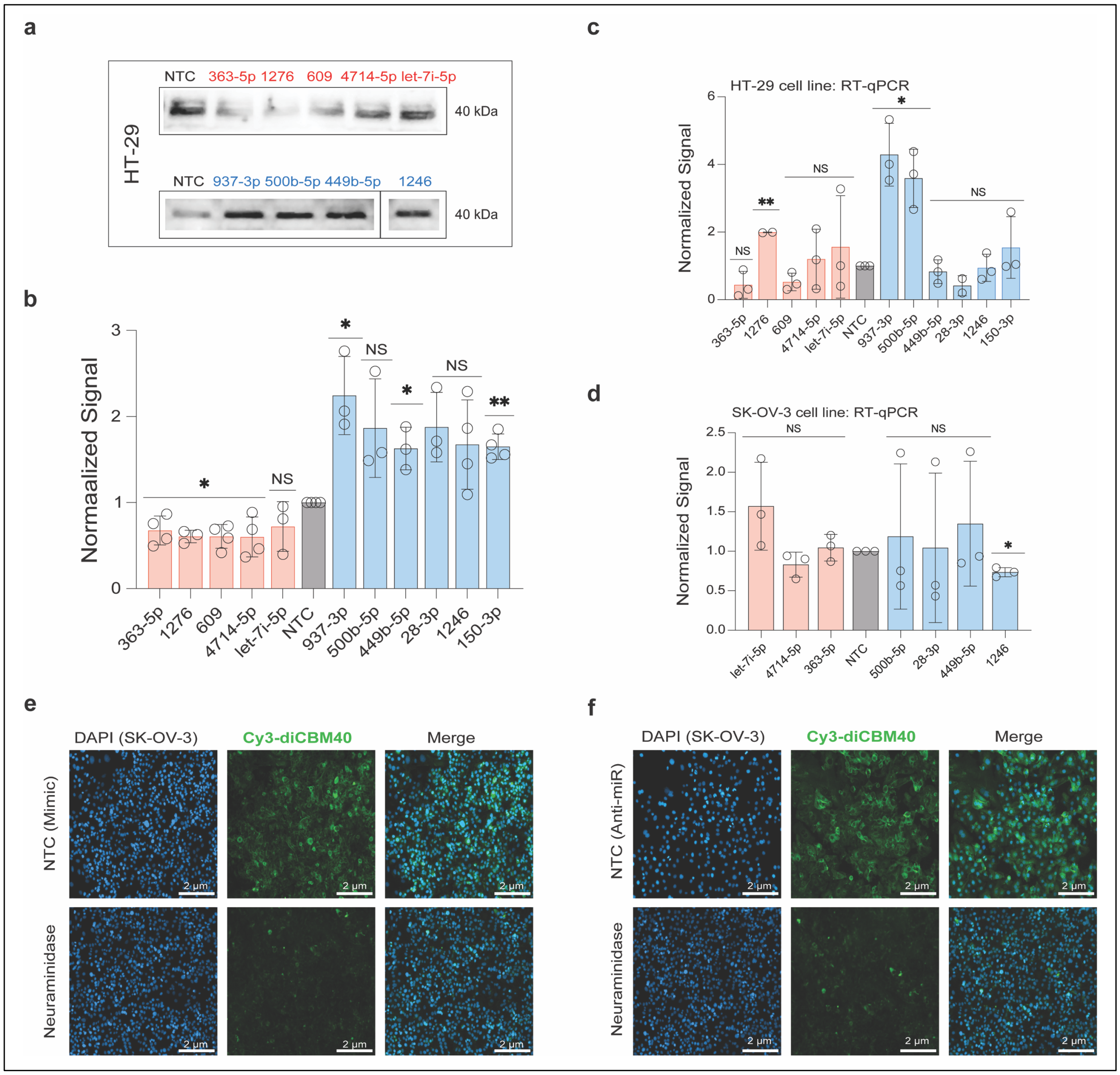


**Supplementary Fig. 11. miRNA up- and downregulate ST3GAL1 expression and α-2,3-sialylation activity in HT-29 and SK-OV-3 cell lines.** Impact of miRNA mimics on ST3GAL1 expression. HT-29 (a, b) cells were treated with miRNA mimics (down-miRs: -363-5p, -1276, -609, let-7i-5p, -4714-5p, up-miRs: -500b-5p, -28-3p, -449b-5p, -937-3p, -150-3p, -1246) or NTC at 50 nM for 48h and analyzed as indicated. **a**. Representative Western blot analysis of ST3GAL1 in HT-29. For simplicity, some wells for upregulators are not shown. Complete blot is in Supplementary Fig. 12h. **b**. Bar graph of Western blot data for HT-29. **c, d**. RT-qPCR quantitative analysis of *st3gal1* in HT-29 (c) and SK-OV-3 (d) cells. **e, f.** Fluorescence microscopy for Cy3-diCBM40 staining in SK-OV-3 cells for miRNA mimic non-targeting control and neuraminidase (e) or anti-miR non-targeting control and neuraminidase (f) treated cells. Additional data are shown in Fig. 4 and Supplementary Figs 10 & 12. All experiments were performed in ≥ biological triplicate. Errors shown are standard deviations. The Wilcoxon *t*-test was used to compare miRs to NTC for Western blot and RT-qPCR experiments (ns not significant, * $p < 0.05$, ** $< 0.01$, *** $< 0.001$).

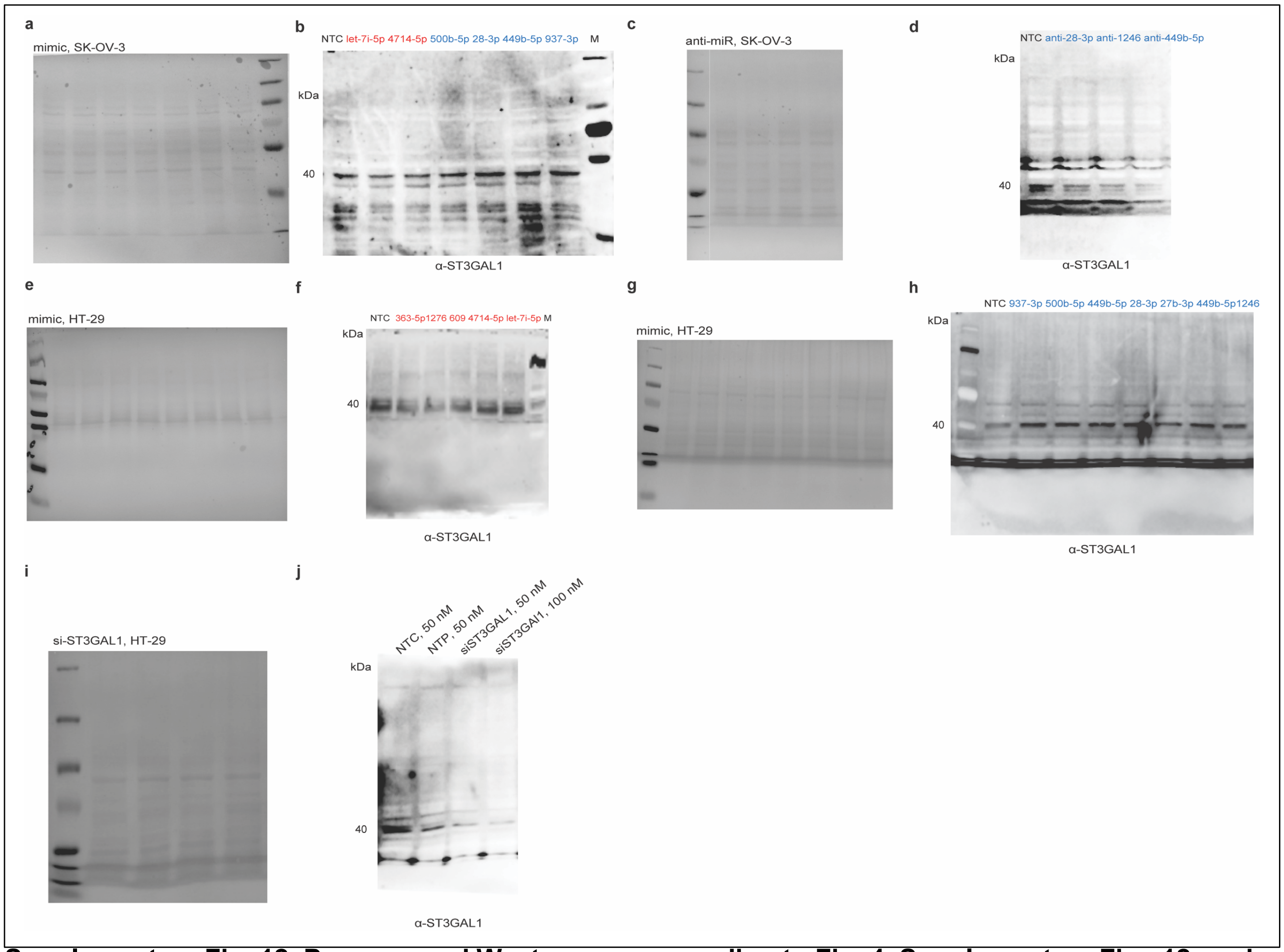


**Supplementary Fig. 12. Ponceau and Western corresponding to Fig. 4, Supplementary Figs 12, and siRNA validation of anti-ST3GAL1 antibody. a, e, g.** Ponceau staining of blot used in Fig. 2 (a: SK-OV-3) and Supplementary Fig. 12 (e & g: HT-29). **b, f, h.** Whole Western blot used in Fig. 4 (b: SK-OV-3) and Supplementary Fig. 11 (f & h: HT-29). **c.** Ponceau staining of blot used in Fig. 4i (c: SK-OV-3) **d.** Whole Western blot for data shown in Fig. 4i (d: SK-OV-3). **i-j.** anti-ST3GAL1 antibody signal validation. **i.** Ponceau staining of blot shown in j (HT-29). **j.** Whole Western blot.

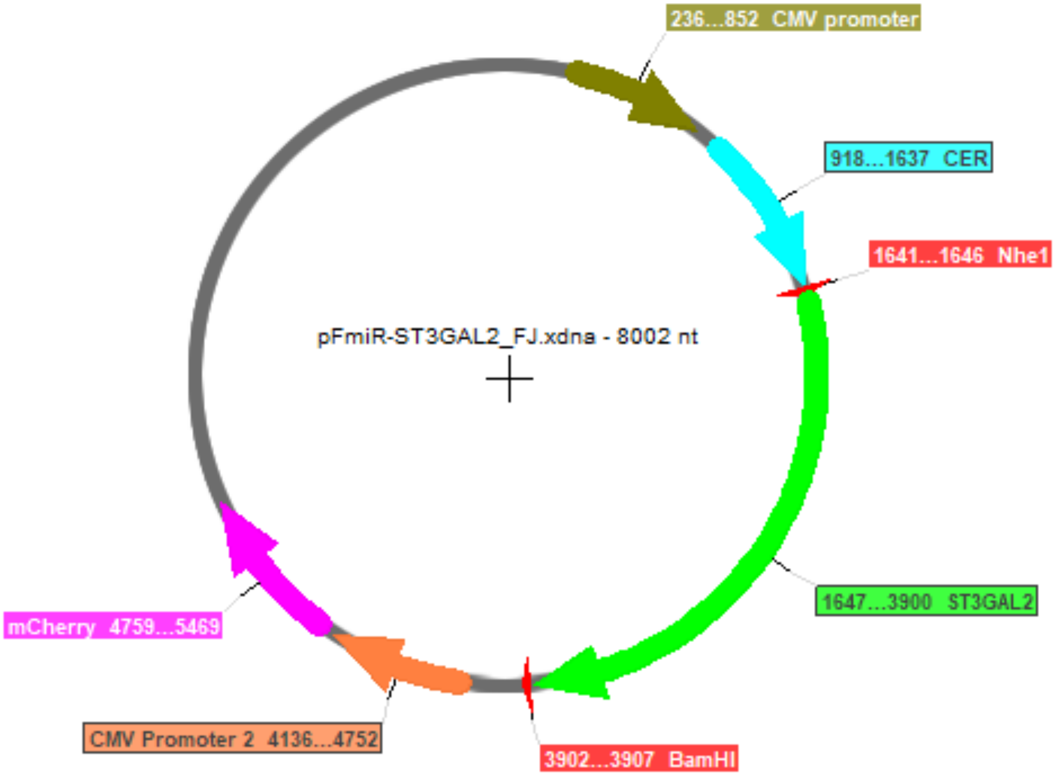


5’GCCGGGCCTCGCCGCGACCCTTCCGGCCCATCTATCGGGCACCGGGGCTCCGGCCCGGGACCCAGGACCAGCAACCCGCGACCAATCATGCTGCAGCCCAGGGGCGTCTGCTGTGCCC
GCCAATCACGAGACTGGGGGACCGGCCGGGCCTGGCACCAATCTGCGCTGCGGTCGGGCGGAGCTTCTGTTTCTCCCAGCCAATCATGTGACTCAAGGAAAACTTCCGGCGCTGTGTCCA
GTCTCCTCCAATCAATGGCCTTCGGGGGCGGGCCAGCGGCCGCTCAATCCCCACTCCCTCATGCTTTGGGTTAGGGTTTTCTTTCACGCTTTCTGAGGAGGAGAGCATGGCGCGGGCCTC
GGCGAAGCACTTCCGTCAGCCTCGGGCGGAGGATCGTCTTAGTAGCTGGCCAGACCTGGAGGAGAGCGGGCGGTCTTTGCGGGCCTCAGGAGATAGGTGGCCGCCCGGCCCCTTTCCCCG
CCTGCCCGGATAGGTGCGTGCAGACCCACCAAAGAAAAGCGGTCGCTGCGGGACCCCCAGTCCCACCTCGGCCGCGCTGTGGGATGGCTGCGCCCCCAGGGCTTCCTGCGTCTCCAGGAA
GCCGGGAGTGGCGCCCTCTGCGGGGTGGGGTGCTCCCGGGCCGAACTCCGGACAGGTGGAGGGGACAGGGCAGAGCTCGAGGAAGCCCTGTCCCCTTCAGTGCAAGGTGCTGTCACTCAC
GTGTGCCCTCGACCCTCCCGTTCACCCGCAGCCTTCTCAGCGCCTCTCCCTGGGCCGGAGGCCTCCTCACCAGCCTACCTGTTGCTCTGGAAAAAAATCCCGTCCCCCGACTCCGTCCCT
ACCCCCAGTCTTCGGCCGGCTCTGGCCCCTGGGGAGGGGGCTGCACGGCGGAAGGAGGCTGGCTATGGGCCCGGCTGCCCGCTGCATGTACCTCCTCCTCCACCCATCGCCTCTTGCCTG
GGGGTAACTTTGCCTGGGGCTCATTCTTTGGTTAAGCTGAAGCTGCCGTGGGTGGCCAAACCGCAGATTCTTTGCAAATTCTGAGCTGGCAGAGCTCGCAGCCGGGAGCCGGCCGGGGAA
GAGGAGACTTGCGCGCCGCAAGCCGCCTGCCTCCACCCTGCTCTCCATCTCCCGCTCTAGAAGGGCTGGGAAGCTCGCGGCCGGGGTTCCACCTGGAAGCTGCTTGCATGGCTGAACCCA
GCTTAGGTCCCTGACGGGGCTGCTGGTGGAATTCTCCCCCTTCGAAGCTGGGGAGGTTTAGGAGGGGGAAGGCTTCTGTGAAGCTCTCAAACCACTAATAGAGCCCCCTCCCCAACAGTG
ACGGCGCAGATGCTCCCCCTTTTCTTAGTTGACACCACCAGGCAGCTTCCTGGCCGTTGGTAGGTTCCTGCAGCTGGCTGAGGGAACAGGGACCGGCAGGGGACTTTGTTAGGGGAGGGT
TGGGATGGGCAGTGGGCCCCTGAAAGTTAATATATTGGAACCTAGCTCGAGTGTCGTTCTTTCCAATTCCGAAAGTAGAAAGAGTAAAAATAGGGGTGATTGGGGTGGGGTTAGTAGAAT
GCCTCTCTCAGGGCGCTCCCCCCTCCCCCACCGTTTTAGAGAGCTAGGCCTCAGCCAGTCTTGCCACTCCCATCTCAGTGCTTCCTGAAGAGGCTGTTTTGAGTGTTGATGAAAAGCAAT
GCAATTATGCCAAACAGTATTGAGCAGAATAATTTATTTCTTTTTTTTTCTTTTGCTTTAAATCATGAATCCCGCCAGGTACGGTGGCTCACACCTGTCATCCCAGCACTTTGGGAGGCCA
AGGCGGGCGGATTACTTAATACTTAAGGTCAGGAGTTCGAGACCAGCCTGGCCAATATGGTGAAACCTCGTCTCTACCAAAAAAAAAAAAAAAATACAAAAATTAGCCAGGCGCAGTGGTG
CGCACCTGTAATCCCAGCTACTTGGAAGGCTGAGGCAGGAGAATCCCTTGAACCGAGGAGGCGAAGGTTGCAGTGAGCCGAGATTGTGCCACTGCACTCCAGCCTGGGCGACAGAGCAAG
ACCCTGTCTCAAAAAAATTAAATCATGAATCCCCATCCTGGAAGAGGTAGGTCCCAGCATCCAGCCAGATTTTCTGCAATAGTAATTTAAACACACTTTTTTATTTCCTTCCCTTTCTGA
ATTAAAAGGAACAAAAAACTGTCTAGTCCTTCATTGTTCTGAAATGGGCATGATGGAAGAAGGCGGGTCTGGTAGGATAGCTGTGACTGATAATG 3’

**Supplementary Fig. 13. pFmiR-ST3GAL2**. **a**. pFmiR-ST3GAL2 plasmid map. **b**. Sequence of ST3GAL2 3'UTR (2254 bp).

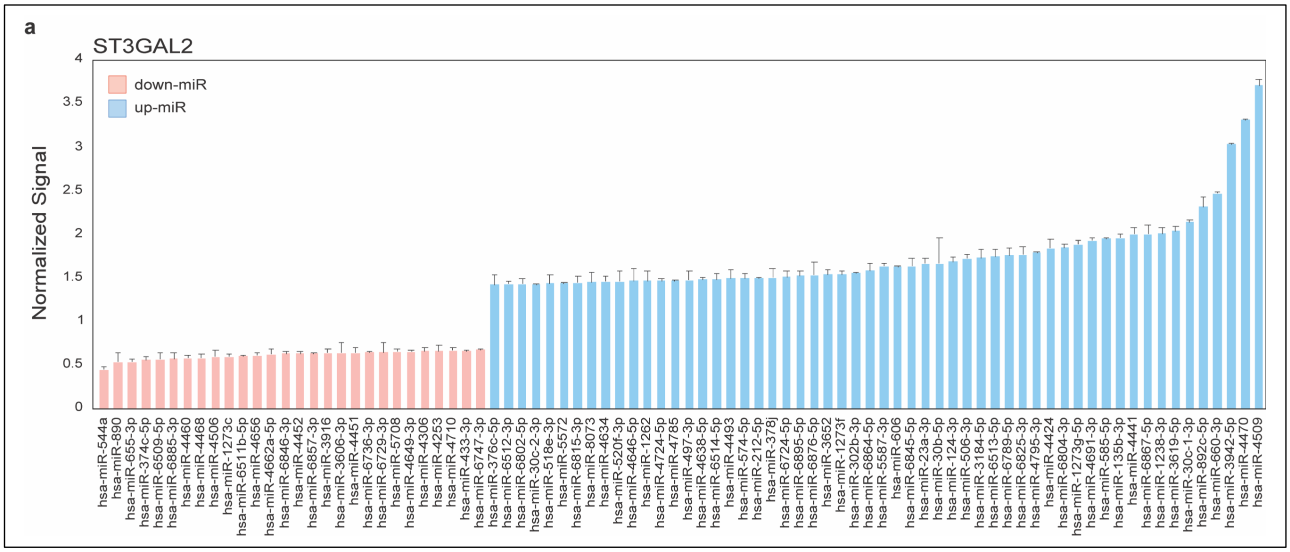


**Supplementary Fig. 14. Hits for ST3GAL2 miRFluR assay**. **a.** Bar graph of miRNA hits for ST3GAL2 (95% confidence interval). Data are normalized over median. Error bars represent standard deviation of technical triplicates.

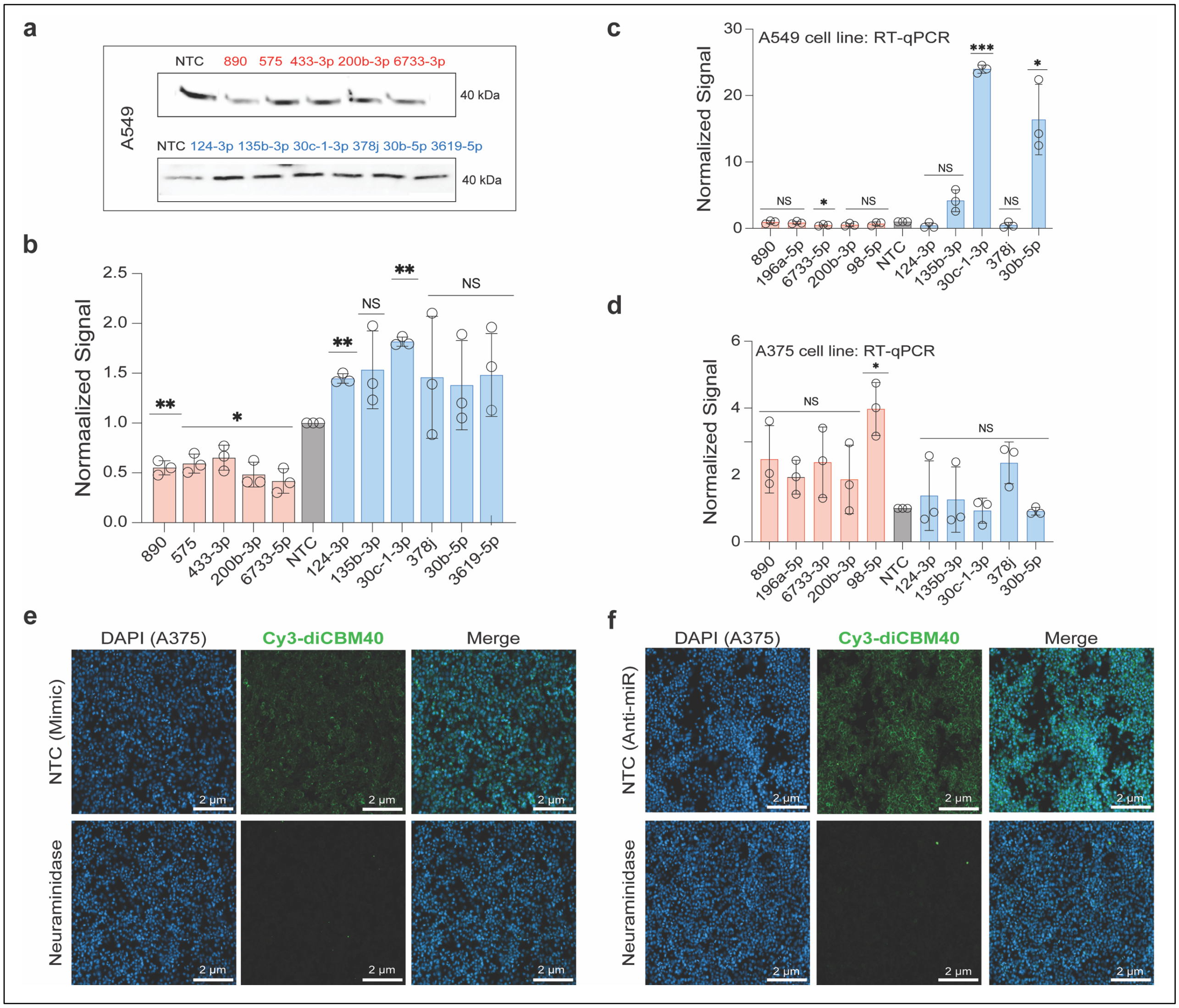


**Supplementary Fig. 15. miRNA up- and downregulate ST3GAL2 expression and α-2,3-sialylation activity in A549 lung and A375 melanoma cell lines.** Impact of miRNA mimics on ST3GAL2 expression. A549 (a, b) cells were treated with miRNA mimics (down-miRs: -890, -575, -433-5p, -200b-3p, -6733-5p, up-miRs: -124-3p, -135b-3p, -30c-1-3p, -378j, -30b-5p, -3619-5p) or NTC at 50 nM for 48h and analyzed as indicated. **a**. Representative Western blot analysis of ST3GAL2 in A549. **e**. Bar graph of Western blot data for A549. **c, d**. RT-qPCR quantitative analysis of *st3gal2* in A549 (c) and A375 (d). **e, f.** Fluorescence microscopy for Cy3-diCBM40 staining on A375 for non-targeting miRNA mimic control and neuraminidase A375 (e) or anti-miR non-targeting control and neuraminidase A375 (f) treated cells. Additional data are shown in Fig. 5 and Supplementary Fig.s 14 & 16. All experiments were performed in ≥ biological triplicate. Errors shown are standard deviations. The Wilcoxon *t*-test was used to compare miRs to NTC for Western blot and RT-qPCR experiments (NS: not significant, * $p < 0.05$, ** $< 0.01$, *** $< 0.001$).

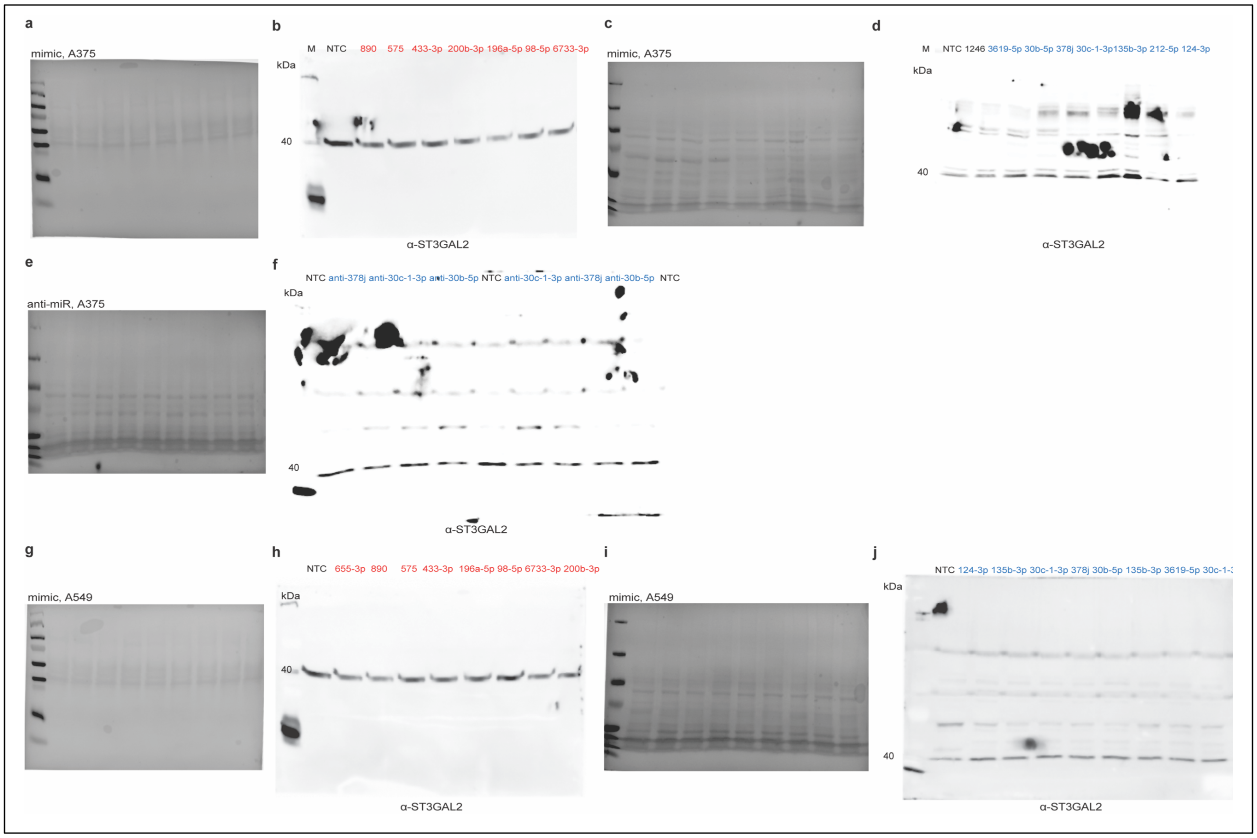


**Supplementary Fig. 16. Ponceau and Western corresponding to Fig. 5, Supplementary Figs 16. a, c, g, i.** Ponceau staining of blot used in Fig. 5 (a & c: A375) and Supplementary Fig. 16 (g & i: A549). **b, d, h, j.** Whole Western blot for data shown in Fig. 5 (b & d: A375) and Supplementary Fig. 16 (h & j: A549). **e.** Ponceau staining of blot shown in Fig. 5i (e: A375) **f**. Whole Western blot data shown in Fig. 5i (f: A375).

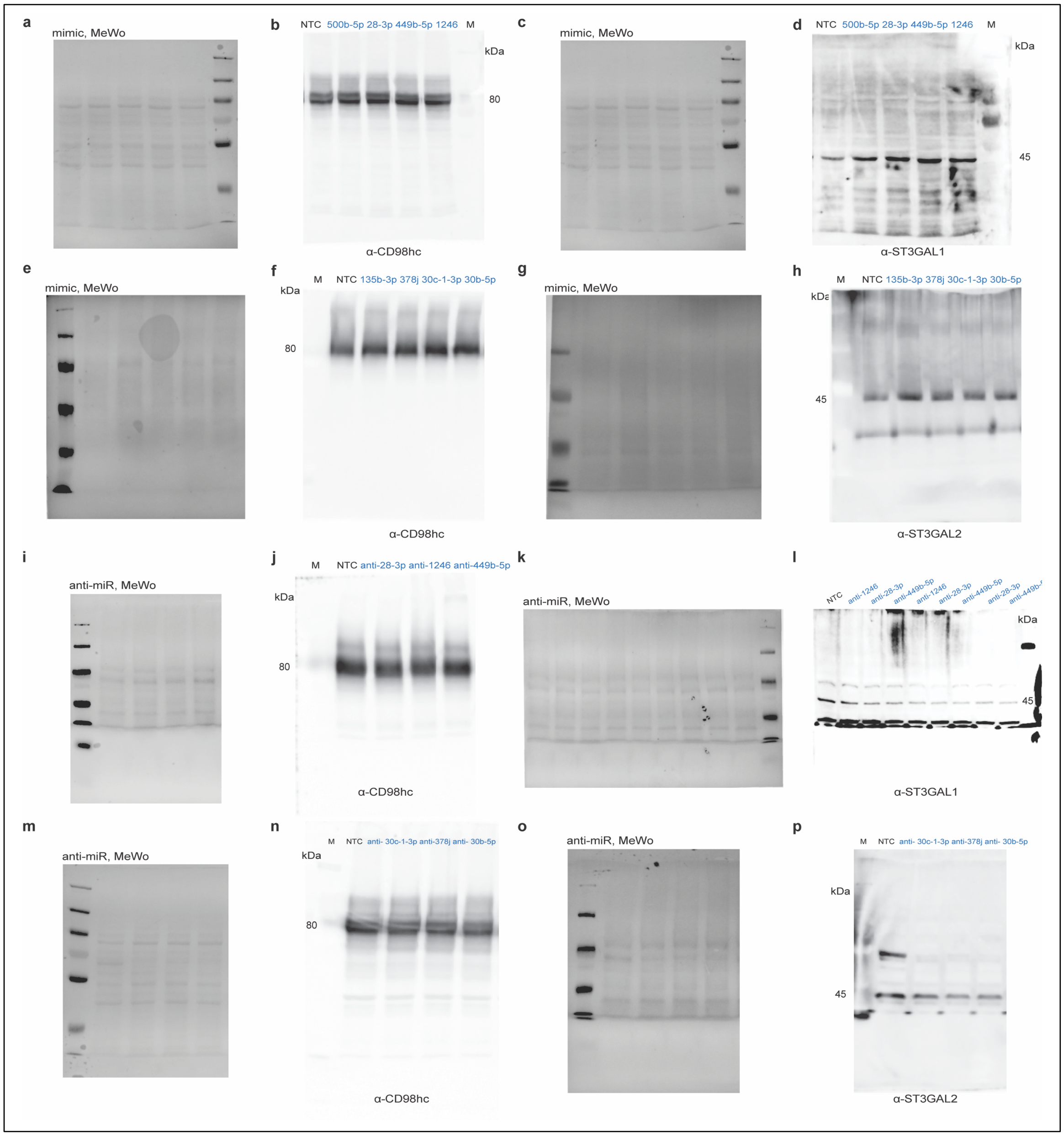


**Supplementary Fig. 17. Ponceau and Western corresponding to Fig. 7. a, c, e, g.** Ponceau staining of blots used in Fig. 7. **b, d, h, j.** Whole Western blots shown in Fig. 7. CD98hc: ST3GAL1 co-upregulation (CD98hc: b; ST3GAL1: d) or CD98hc: ST3GAL2 upregulation (CD98hc: f; ST3GAL2: h)**. i, k, m, o.** Ponceau staining of blots used in Fig. 7. **j, l, n, p.** Whole Western blots for data shown in Fig. 7. CD98hc: ST3GAL1 co-upregulation (CD98hc: j; ST3GAL1: l) or CD98hc: ST3GAL2 upregulation (CD98hc: n; ST3GAL2: p)**.**

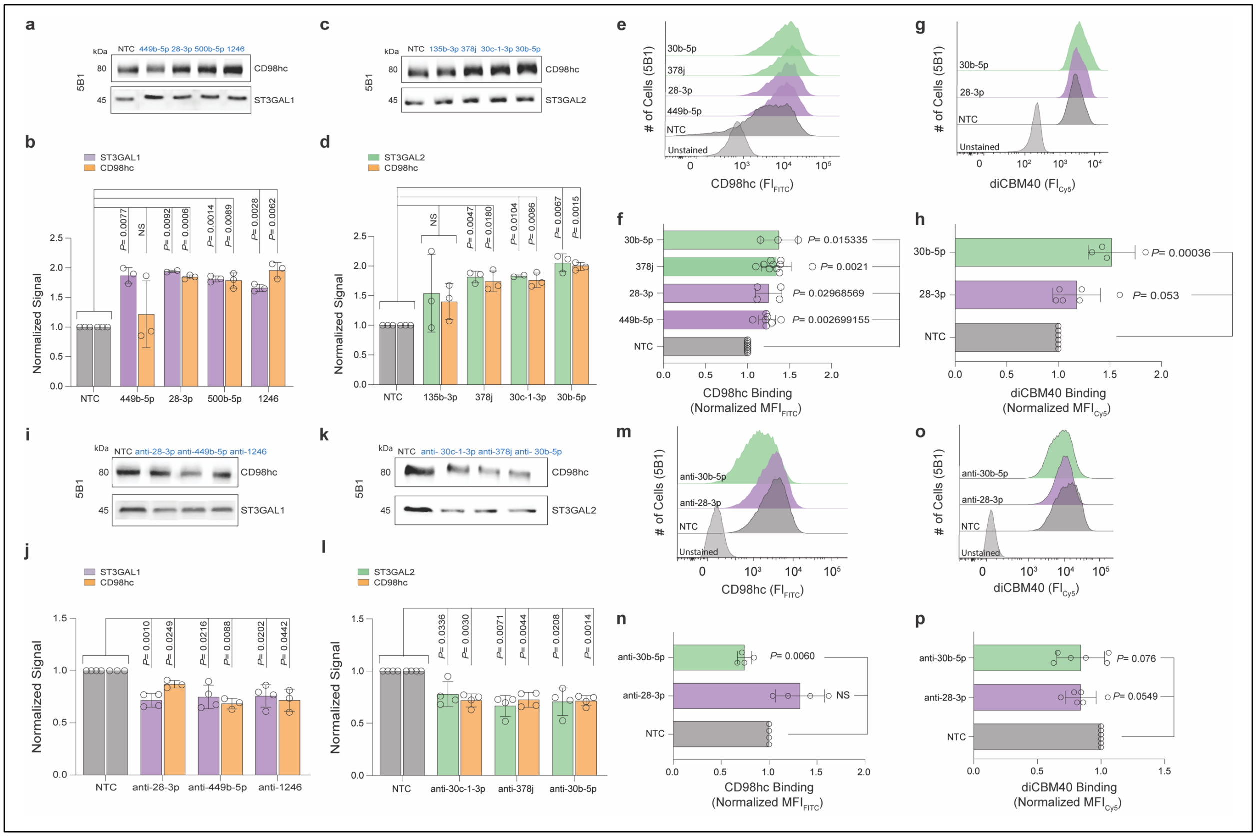

**Supplementary Fig. 18. Co-upregulation of CD98hc and either ST3GAL1 or ST3GAL2 is observed in melanoma. a-d.** Impact of co-up-miR mimics on either CD98hc (orange) & ST3GAL1 (purple, co-up-miRs: -500b-5p, -28-3p, -449b-5p, -1246) or CD98hc (orange) & ST3GAL2 expression (green, co-up-miRs: -135b-3p, -378j, -30c-1-3p, -30b-5p) in 5B1 cells treated with corresponding co-up-miRs or non-targeting control (NTC) at 50 nM for 48h and analyzed as indicated. **a**. Representative Western blot analysis of CD98hc and ST3GAL1. **b**. Bar graph of corresponding Western blot data. **c**. Representative Western blot analysis of CD98hc and ST3GAL2. **d**. Bar graph of corresponding Western blot data. **e**. Representative flow cytometry analysis of CD98hc binding. **f**. Bar graph of flow cytometry data. **g**. Representative flow cytometry analysis of Cy5-diCBM40 binding. **h**. Bar graph of flow cytometry data. **i-o.** Impact of endogenous co-up-miRs on either CD98hc (orange) & ST3GAL1 (purple) or CD98hc (orange) & ST3GAL2 (green) expression. 5B1 cells were treated with anti-miRs (anti-up-miRs: anti-28-3p, anti-1246, anti-449b-5p; anti-30c-1-3p, anti-378j, anti-30b-5p) or non-targeting control (NTC) at 50 nM for 48h and analyzed as indicated. **i**. Representative Western blot analysis of CD98hc and ST3GAL1. **j**. Bar graph of Western blot data. **k**. Representative Western blot analysis of CD98hc and ST3GAL2. **l**. Bar graph of Western blot data. **m.** Representative flow cytometry analysis of CD98hc binding. **n.** Bar graph of flow cytometry data for 5B1. **o.** Representative flow cytometry analysis of Cy5-diCBM40 binding. **p.** Bar graph of flow cytometry data. Additional data are shown in Fig.7 and Supplementary Fig.s 17, 19-21. All experiments were performed in ≥ biological triplicate. Errors shown are standard deviations. For Western blot analysis, the Wilcoxon *t*-test was used to compare miRs to NTC (NS: not significant, $^{*}p < 0.05$, $^{**} < 0.01$, $^{***} < 0.001$). For flow cytometry analysis, paired *t*-test was used to compare miRs to NTC and *p*-values are indicated on the graph.

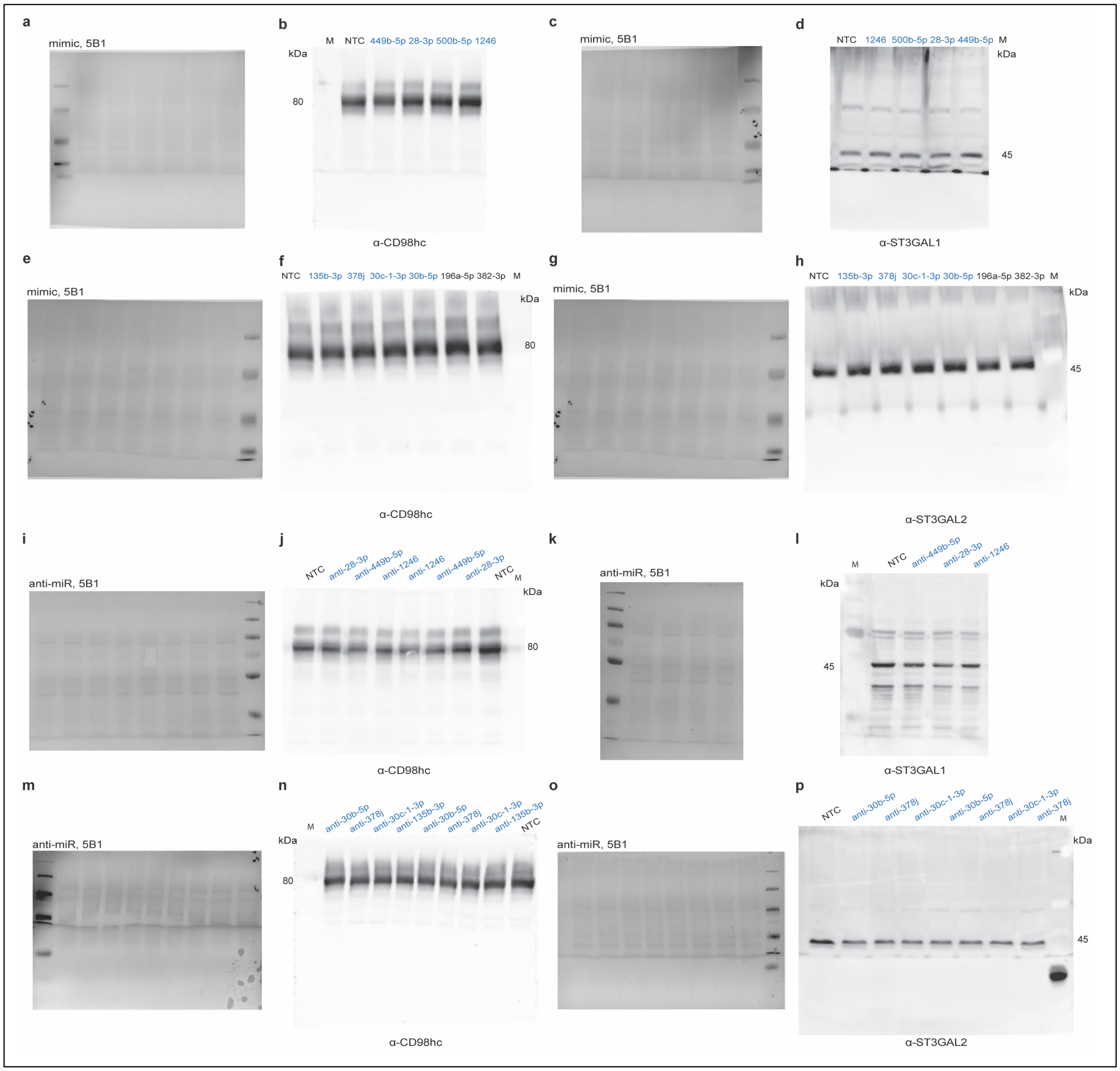


**Supplementary Fig. 19. Ponceau and Western corresponding to Supplementary Fig. 19. a, c, e, g.** Ponceau staining of blots used in Supplementary Fig. 19. **b, d, h, j.** Whole Western blots for data shown in Supplementary Fig. 19. CD98hc: ST3GAL1 co-upregulation (CD98hc: b; ST3GAL1: d) or CD98hc: ST3GAL2 upregulation (CD98hc: f; ST3GAL2: h)**. i, k, m, o.** Ponceau staining of blots shown in Supplementary Fig. 19. **j, l, n, p.** Whole Western blots for data shown in Supplementary Fig. 19. CD98hc: ST3GAL1 co-upregulation (CD98hc: j; ST3GAL1: l) or CD98hc: ST3GAL2 upregulation (CD98hc: n; ST3GAL2: p)**.**

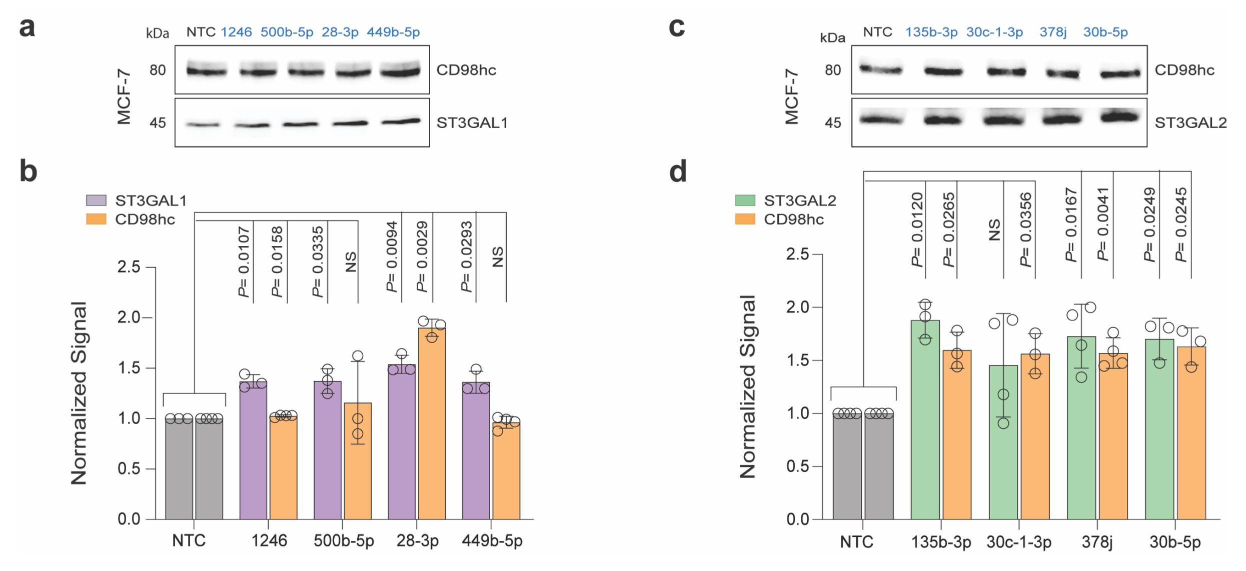


**Supplementary Fig. 20. miRNA regulation of CD98hc and α-2,3-sialylation in MCF-7 cell line.** Impact of co-up-miR mimics on either CD98hc (orange) & ST3GAL1 (purple, co-up-miRs: -500b-5p, -28-3p, -449b-5p, -1246) or CD98hc (orange) & ST3GAL2 expression (green, co-up-miRs: -135b-3p, -378j, -30c-1-3p, -30b-5p) in MCF-7 cells treated with corresponding co-up-miRs or non-targeting control (NTC) at 50 nM for 48h and analyzed as indicated. **a**. Representative Western blot analysis of CD98hc and ST3GAL1. **b**. Bar graph of Western blot data for blots shown in a. **c**. Representative Western blot analysis of CD98hc and ST3GAL2. **d**. Bar graph of Western blot data for blots shown in c. Additional data are shown in Fig. 7 and Supplementary Fig.s 17-19 & 21. All experiments were performed in ≥ biological triplicate. Errors shown are standard deviations. For Western blot analysis the Wilcoxon *t*-test was used to compare miRs to NTC (NS: not significant, * *p < 0.05,* ** *< 0.01,* *** *< 0.001*). In flow cytometry analysis, paired *t* test was used to compare miRs to NTC and *p*-values are indicated on the graph.

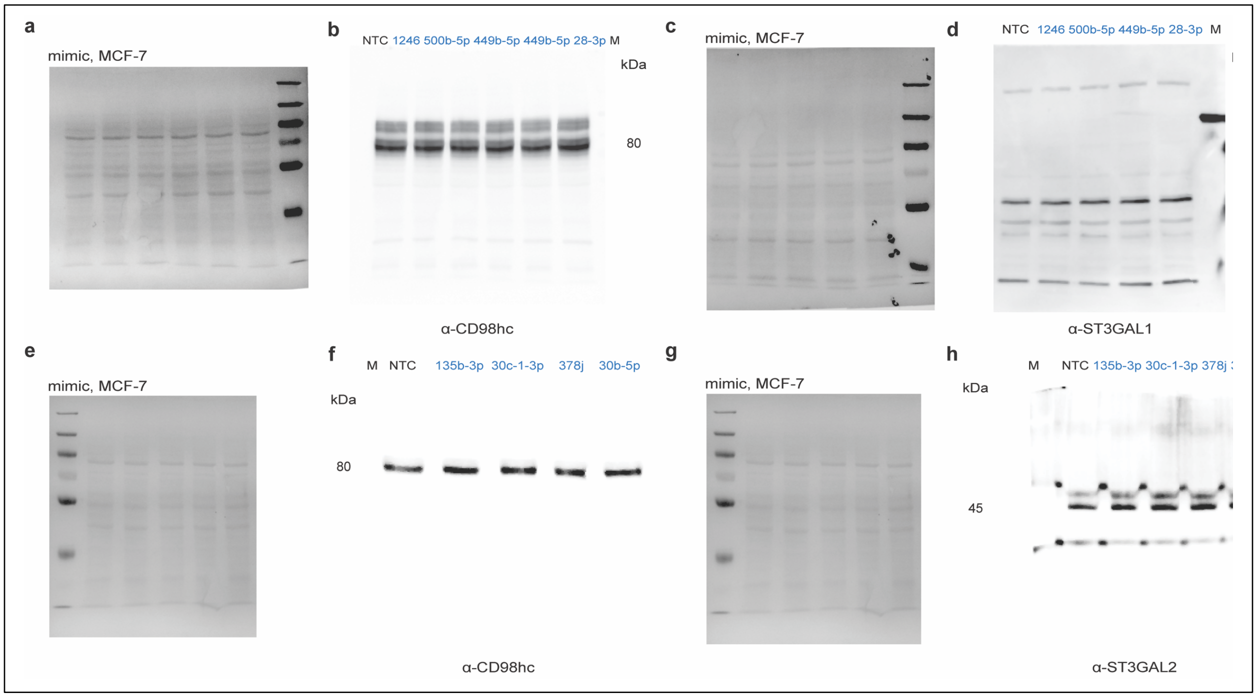


**Supplementary Fig. 21. Ponceau and Western corresponding to Supplementary Fig. 20. a, c, e, g.** Ponceau staining of blots used in Supplementary Fig. 21. **b, d, f, h.** Whole Western blots for data shown in Supplementary Fig. 21. CD98hc: ST3GAL1 co-upregulation (CD98hc: b; ST3GAL1: d) or CD98hc: ST3GAL2 upregulation (CD98hc: f; ST3GAL2: h).

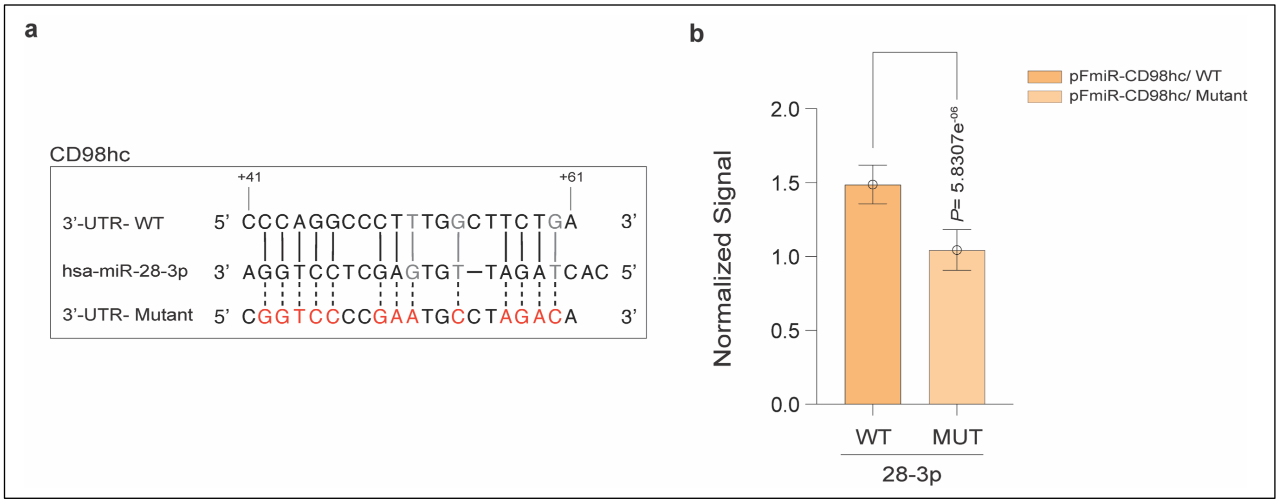


**Supplementary Fig. 22. Mutational analysis identifies miR-28-3p interaction site in CD98hc 3'UTR. a**. Alignment of miR-28-3p with predicted CD98hc-3′UTR site and its corresponding mutant. Mutated residues are shown in red, Wobble interactions (G:U) in grey. **b**. Bar graph of data from mutant miRFluR sensor as in b. Data were normalized over NTC in each sensor. Statistical analysis using the standard *t*-test compared the impact of miR-28-3p in the WT sensor with the corresponding mutant.

**Supplementary Table 1.** CD98hc up-miR analysis in melanoma dataset. Co-up-miRs with ST3GAL1- *purple,* ST3GAL2- *green*. (Gómez-Martínez, *et. al.*), (*Gaziel-Sovran, *et. al.*).

| up-miR | fold change in miRFluR (normalization over control) | signature in melanoma (Log fold-change > 0) |
|---|---|---|
| hsa-miR-4640-5p | 1.289747317 | up |
| hsa-miR-1246 | 1.299015856 | up |
| hsa-miR-135b-5p | 1.303684609 | up |
| hsa-miR-4436b-3p | 1.321963315 | up |
| hsa-miR-30b-5p* | 1.326657787 | up |
| hsa-miR-30c-1-3p | 1.357529203 | up |
| hsa-miR-6784-5p | 1.380467402 | up |
| hsa-miR-34c-3p | 1.407505853 | up |
| hsa-miR-199a-3p | 1.44020141 | up |
| hsa-miR-660-5p | 1.467075023 | up |
| hsa-miR-135b-3p | 1.487461446 | up |
| hsa-miR-500a-3p | 1.489550747 | up |
| hsa-miR-299-3p | 1.554398492 | up |
| hsa-miR-28-3p | 1.556006354 | up |
| hsa-miR-155-5p | 1.652779561 | up |

**Supplementary Table 2.** Primer sequences for PCR amplification of WT 3'UTRs (a), RT-qPCR quantification of mRNAs (b) Site Directed Mutagenesis (c) for CD98hc, ST3GAL1 and ST3GAL2.

| Primer Name | Sequence (5' → 3') | Sample |
|---|---|---|
| a. PCR amplification of CD98hc, ST3GAL1 and ST3GAL2 3'UTRs | | |
| CD98hc-FWD[a] | AGGTAGCTAGCCTTCAGCCTGACATGGAC | gDNA, MCF-7 |
| CD98hc-REV[a] | AGGTAGGATCCTATGAGAGAAGCAGAGGGAA | gDNA, MCF-7 |
| ST36GAL1-FWD | AGTCAACCGGTAGATGACGCAGTGAAGGG | gDNA, MCF-7 |
| ST3GAL1-REV | AGTCACGTACGAACAATAAAATAGCTCTTTGTTTATTCAC | gDNA, MCF-7 |
| ST3GAL2-FWD | AGTCAGCTAGCAAGTCTACCGGGGCAACTGAG | gDNA, MCF-7 |
| ST3GAL2-REV | AGTCAGGATCCCATTATCAGTCACAGCTATCCTACC | gDNA, MCF-7 |
| b. RT-qPCR[a] quantification of CD98hc, ST3GAL1, ST3GAL2 or GAPDH transcript | | |
| ST3GAL1-FWD | TTGGAGGACGACACCTACCGAT | Total RNA, HT-29, SK-OV-3 |
| ST3GAL1-REV | CACCACTCTGAACAGCTCCTTG | Total RNA, HT-29, SK-OV-3 |
| ST3GAL2-FWD | TCCGACTGGTTTGACAGCCACT | Total RNA, A549, A375 |
| ST3GAL2-REV | CTTCTCCAGCACCTCATTGGTG | Total RNA, A549, A375 |
| CD98hc-FWD | CCCACTACCCTTCTCCTTTCCTT | Total RNA, 5B1, MCF-7 |
| CD98hc-REV | GTTCACTCATAATCTGCAACAGTTTG | Total RNA, 5B1, MCF-7 |
| GAPDH-FWD | GGTGTGAACCATGAGAAGTATGA | Total RNA, all cell lines |
| GAPDH-REV | GAGTCCTTCCACGATACCAAAG | Total RNA, all cell lines |
| c. PCR amplification of CD98hc, ST3GAL1 and ST3GAL2 mutant 3'UTRs | | |
| 155-5p-CD98hc-MUT-FWD | accagatcacTCCCTCTGCTTCTCTCATAC | pFmiR-CD98hc |
| 155-5p-CD98hc-MUT-REV | acgtccccagACTTTTATTTGAAGGCAGAAAAC | pFmiR-CD98hc |
| 28-3p-CD98hc-MUT-FWD | tggctagacaTTTTTCTCTTTTTTAAAAACAAACAAAC | pFmiR-CD98hc |
| 28-3p-CD98hc-MUT-REV | ttcggggaccgAAGGAAAGGAGAAGGGTAG | pFmiR-CD98hc |
| 30b-5p-CD98hc-MUT-FWD | cccaaaatcccacaaatCTGCCTTCAAATAAAAGTCAC | pFmiR-CD98hc |
| 30b-5p-CD98hc-MUT-REV | ggttgtgagttaatctgCAACAGTTTGTTTGTTTGTTTTTAAAAAAG | pFmiR-CD98hc |
| 1246-CD98hc-MUT-FWD | tcttcagatCCCTCTGCTTCTCTCATAC | pFmiR-CD98hc |
| 1246-CD98hc-MUT-REV | ccatcgtccgGTGACTTTTATTTGAAGGCAG | pFmiR-CD98hc |
| 1246-ST3GAL1-MUT-FWD | ttttttaggtCCAAAACTTTAAAGACTTTTCTTTTTTC | pFmiR-ST3GAL1 |
| 1246-ST3GAL1-MUT-REV | gcactttgtcctTTTCGGTTTGTTTAACAC | pFmiR-ST3GAL1 |
| 30b-5p-ST3GAL2-MUT-FWD | cctacgcacgccCCCCAGGGCTTCCTGCGT | pFmiR-ST3GAL2 |
| 30b-5p-ST3GAL2-MUT-REV | gtgtgggggcgggAGGTGGGACTGGGGGTCC | pFmiR-ST3GAL2 |

[a] FWD, forward; REV, reverse; RT-qPCR, Reverse transcription quantitative polymerase chain reaction.

**Dataset 1-3. (separate excel file) | miRFluR results for CD98hc (Dataset 1), ST3GAL1 (Dataset 2) and ST3GAL2 (Dataset 3)**. Data analysis before and after thresholds.
